# Evaluation of deep learning tools for chromatin contact prediction

**DOI:** 10.64898/2026.02.27.708323

**Authors:** Hoang Thien Thu Nguyen, Vanessa Vermeirssen

## Abstract

**Background:** Three-dimensional chromatin organization plays a central role in gene regulation and is commonly measured using Hi-C technology. Recently, deep learning models have been developed to predict Hi-C contact maps from genomic and epigenomic features, offering a computational alternative to costly experimental assays. However, the performance, robustness, and biological interpretability of these models remain unclear due to the absence of systematic benchmarking.

**Results:** We present a comprehensive benchmark to evaluate five Hi-C prediction models, C.Origami, Epiphany, ChromaFold, HiCDiffusion, and GRACHIP, across multiple evaluation criteria, including accuracy, visual fidelity and loop detection. Among all models, Epiphany achieved the strongest overall performance, combining high accuracy, cell-type generalization, realistic image quality and reliable loop detection. Moreover, we evaluated predicted contact maps using four different loop-callers to assess the impact of model choice on loop detection performance. Despite the coarse resolution, many models could recover biologically relevant interactions. Notably, structural map quality was more critical than the choice of loop-caller for reliable detection. Finally, ablation analyses revealed that epigenomic signals are influential features for accurate Hi-C prediction. Despite the use of multiple input modalities in many models, only a limited subset contributed substantially to predictive performance.

**Conclusions:** This study provides a systematic comparison of deep learning models for Hi-C prediction and highlights the importance of specific regulatory signals in reconstructing 3D chromatin organization. The proposed evaluation framework clarifies model behaviours and offers guidance for the development and interpretation of Hi-C prediction methods.

## Introduction

Gene expression is tightly controlled by a complex network of regulatory mechanisms that ensure genes are expressed at the right time, location, and level [1]. Among the various layers of regulation, transcriptional regulation plays a central role [2]. At this level, transcription factors (TFs) bind to promoter regions and to distal regulatory elements such as enhancers to modulate gene activity. Because enhancers can act over long genomic distances and regulatory interactions are often combinatorial, the architecture of gene regulatory networks (GRNs) is highly complex, dynamic, and context-dependent. A key source of complexity arises from the fact that many genes are controlled by distal enhancers, which may be located tens to hundreds of kilobases, or even megabases, away from their target promoters. Early studies often assumed that an enhancer regulates its nearest gene along the linear genome sequence [3]. However, this distance-based assumption has proven to be overly simplistic because the genome is folded in three-dimensional (3D) space within the nucleus, allowing distal regulatory elements to physically contact promoters [4]. Accordingly, enhancers defined from chromatin accessibility data frequently act over long genomic distances: 80.94% are located more than 2 kb away from the nearest transcription start site [5]. Moreover, these distal enhancers tend to be less conserved and more cell type-specific and hence are crucial for gene regulation biology. For instance, ZRS enhancer is located in the intron of *LMBR1*, around 1 Mb away from *SHH*, which controls limb expression [6]. Enhancer variants within the *FTO* locus were shown to act through long-range chromatin interactions to regulate *IRX3/IRX5* rather than the nearest gene [7].

To capture these long-range enhancer-gene interactions, researchers have developed technologies to decipher 3D chromatin contacts, among which Hi-C has become the most widely used [8]. Hi-C is a genome-wide chromosome conformation capture (3C) technique in which chromatin is crosslinked, restriction-digested, and proximity-ligated, followed by paired-end sequencing to identify genomic loci that are physically close in the nucleus. Hi-C revealed that 3D chromatin organization is hierarchical. At the megabase scale, chromosomes segregate into A and B compartments corresponding to active and inactive chromatin, respectively. At an intermediate scale, the genome is partitioned into topologically associating domains (TADs), regions with enriched internal interactions. At a finer scale, chromatin loops connect specific loci, often linking enhancers and promoters. Chromatin looping within TADs constrains enhancer–promoter interactions and contributes to cell-type-specific gene regulation [8]. One of the major advantages of Hi-C is its scalability and high throughput, allowing the detection of fine-scale chromatin interactions such as enhancer-promoter loops [9, 10]. Furthermore, Hi-C data can be integrated with other omics datasets to provide a comprehensive view of gene regulation. However, mapping 3D genome interactions with Hi-C remains technically challenging and prohibitively costly. High-resolution chromatin contact maps (e.g., ∼5 kb resolution requiring ∼2 billion read pairs) are therefore available for only a limited number of cell types and experimental conditions. To overcome the scarcity of 3D chromatin organization data, researchers have increasingly turned to computational prediction methods, particularly deep learning approaches. These models aim to infer chromatin interactions or Hi-C contact maps using multi-omics data. In computational terms, this task is often described as modality prediction, where one or more data modalities are used to predict another modality. This strategy reduces experimental costs and enables reconstruction of missing biological layers that is otherwise experimentally difficult to obtain. In sequence-to-function models, regulatory information is directly learnt from genomic DNA sequence to predict diverse molecular readouts such as chromatin accessibility, transcription factor binding, and gene expression, thereby linking genotype to gene regulatory function [11]. Several successful examples of modality prediction have been reported. For instance, BABEL predicts chromatin accessibility from gene expression, while MultiVI can impute missing RNA or ATAC profiles in partially observed single-cell multiome datasets [12, 13].

Recent years have seen a rapid expansion of deep learning-based models predicting 3D chromatin contact organization from DNA sequence and commonly generated epigenomic tracks. Convolutional neural network (CNN)-based models have been particularly successful at predicting genome-wide chromatin interactions [14], while transformer-based models have further advanced the field by capturing long-range dependencies through self-attention mechanisms [15]. Hybrid architectures that combine CNNs and transformers have shown additional improvements by leveraging the strengths of both, extracting local sequence features with CNNs and modelling long-range relationships with transformers [16]. The development of these models has significantly improved our ability to predict Hi-C maps from accessible data sources such as Assay for Transposase-Accessible Chromatin using sequencing (ATAC-seq) and Chromatin Immunoprecipitation of CCCTC-binding factor (CTCF ChIP / CUT & RUN). Various other architectures, including some generative models, have been applied to this problem too [17]. These approaches not only aim to maximize predictive accuracy but also to enhance the quality, generalizability, and biological interpretability of predicted Hi-C maps. However, 3D chromatin architecture is strongly cell-type specific. Models relying only on DNA sequence lack regulatory context and therefore tend to predict similar structures across cell types. This limitation has motivated the development of multimodal and context-aware models that incorporate additional molecular measurements. The models compared in this study differ substantially in their input modalities and architectural assumptions. Epiphany predicts Hi-C contact maps using only epigenomic tracks, such as chromatin accessibility and histone modifications, combining with generative adversarial networks (GAN) model, enabling cell-type-specific prediction and visual fidelity [18]. ChromaFold instead focuses on using single-cell ATAC-seq (scATAC-seq) accessibility profiles, allowing recovery of cell-type-specific chromatin interactions from a few number of modalities [19]. GRACHIP integrates DNA sequence and genomic features within a graph neural network framework to explicitly model relationships between interacting loci [20]. HiCDiffusion predicts chromatin interactions directly from genomic DNA sequence only using a transformer encoder combined with a diffusion-based generative module. This model primarily investigates whether DNA sequence alone can sufficiently capture cell-type-specific Hi-C contact maps [17]. In contrast, C.Origami, which is often used as a benchmark model, primarily learns general chromatin folding patterns from sequence-derived features and does not inherently model cell-type-specific regulatory variation [21]. These approaches aim not only to improve prediction accuracy but also to reconstruct biologically meaningful chromatin features, enabling inference of genome organization from widely available multi-omics data such as ATAC-seq and CTCF profiling. However, a major challenge remains: with many deep learning models now available, which one performs best, and under what criteria? Previous models were typically evaluated only against a limited set of baseline models (e.g., C.Origami [21] or Akita [22]) in their own method paper, focusing primarily on accuracy while neglecting other aspects such as generalization and biological relevance. Consequently, there is no standardized framework, nor neutral benchmarking effort, for evaluating or comparing deep learning models for Hi-C prediction, making it difficult to assess their relative strengths and weaknesses. A comprehensive benchmarking effort must therefore consider multiple evaluation dimensions, including robustness across cell types, and the biological plausibility of predicted structures.

In this manuscript, we aim to systematically compare five representative models: C.Origami [21], Epiphany [18], ChromaFold [19], HiCDiffusion [17], and GRACHIP [20]. The evaluation focussed on multiple perspectives, including predictive accuracy, visual quality of the generated maps, generalization ability across different cell types or species, and biological relevance for downstream analyses. In addition, we performed ablation analyses to assess the necessity and contribution of each input modality to model performance. These criteria were used to determine how closely the predicted Hi-C maps resemble experimental data and how effectively each model captured meaningful chromatin interactions. Through this neutral benchmarking, we identified the architectural designs, input features, and training strategies that contribute to superior model performance. Ultimately, this framework establishes a reference for standardized evaluation in future studies of 3D genome modelling and promotes the reliable use of in silico Hi-C prediction in downstream applications.

## Results

We compared five representative models for 3D chromatin contact prediction - C.Origami, Epiphany, ChromaFold, HiCDiffusion, and GRACHIP, using the evaluation workflow summarized in Figure 1. For each model, input data were pre-processed according to the recommended settings, and predicted Hi-C contact maps were generated across four human cell lines and 22 chromosomes. The predicted maps were then compared with experimental Hi-C data to assess model performance from multiple complementary perspectives. The predictions generated by each model were assessed using three criteria: (1) predictive accuracy, (2) image quality, and (3) loop detection performance. Predictive performance was firstly evaluated by direct comparison of predicted and experimental contact matrices using three quantitative metrics: mean squared error (MSE), insulation score correlation, and observed-over-expected (O/E) Hi-C correlation. To further examine the contribution of input information, ablation analyses were performed by removing individual modalities and measuring the resulting performance changes. Second, we evaluated the visual realism of predicted contact maps using the Fréchet Inception Distance (FID). Finally, predicted contact maps were processed with a loop-calling pipeline, and the detected chromatin loops were compared with loops identified from experimental Hi-C data to assess downstream biological utility.

**Figure 1.**
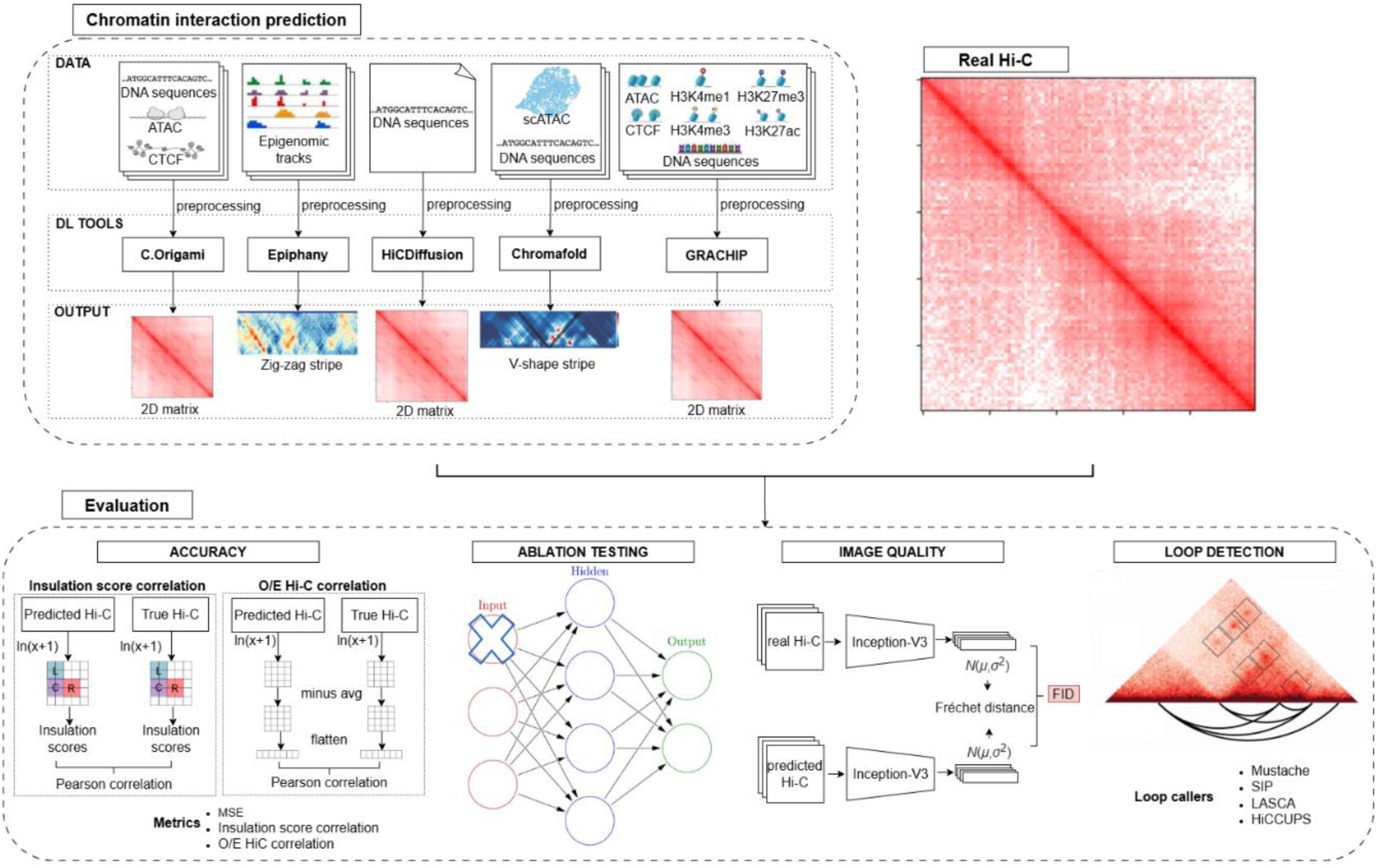
Overview of this Hi-C prediction benchmarking study. Input data, including DNA sequences and various epigenomic features, were first preprocessed according to the protocols described in the original studies for each model. The processed data were then used to train multiple deep learning models (C.Origami, Epiphany, HiCDiffusion, Chromafold, and GRACHIP), following their respective training procedures and dataset configurations. Each trained model was used to predict Hi-C contact maps. Both the predicted and real Hi-C data were further preprocessed to ensure a comparable format. Model performance was evaluated using several metrics. For accuracy, we computed MSE, insulation score correlation, and O/E Hi-C correlation. Insulation score correlation was calculated by first applying a natural logarithm transformation to the Hi-C matrices, then computing insulation scores, and finally measuring the Pearson correlation between the predicted and ground-truth insulation profiles. For O/E Hi-C correlation, each contact matrix was normalized by subtracting its mean to account for scale differences, then flattened into a vector, and the Pearson correlation was computed between predicted and real matrices. Ablation testing was conducted to assess the contribution of different input modalities. Image quality was evaluated using Fréchet Inception Distance (FID), where features were extracted using an InceptionV3 network and the Fréchet distance was computed between the feature distributions of predicted and real Hi-C images. Finally, loop detection analysis was performed using four loop-calling algorithms: Mustache, SIP, LASCA, and HiCCUPS.

### Accuracy performance evaluation

In this section, we compared the predictive accuracy of each model across different cell lines and performed a cross-model benchmark to identify the best-performing approach. Model performance was assessed using the quantitative performance metric MSE and the more biologically relevant metrics (insulation score correlation and O/E Hi-C correlation) (Methods). The insulation score correlation measures how well a model captures local chromatin domain structure by comparing insulation profiles, whereas O/E Hi-C correlation measures similarity of 3D contact patterns after correcting for distance-dependent contact decay.

#### a. Within-model generalization across cell types and chromosomes

First, each model was evaluated independently by examining its performance by MSE scoring across different cell lines and chromosomes (Figure 2). The top panel showed a bubble plot summarizing model performance across chromosomes and cell lines, while the bottom panel displayed violin plots illustrating the distribution of scores. Three notable patterns were observed. First, an unexpected pattern was observed for HiCDiffusion, ChromaFold, and GRACHIP, where performance on the training cell line was lower than on unseen cell lines. Second, the variability of MSE scores was very large, with training performance ranging from near 0 to near 1 depending on the chromosome and cell line. Third, C.Origami showed very high performance on the training data but extremely low values on unseen cell lines, with scores close to 0, suggesting a complete failure of prediction (C.Origami). Although previous reports that C.Origami was known to have limited ability to capture cell-type-specific regulatory variation, performance values near zero are unexpected, as they would imply a complete loss of predictive signal. Therefore, the near-zero scores cannot be explained solely by overfitting, as even an overfitted model would be expected to retain some predictive signal. Instead, this observation suggests that the MSE-based metric may not accurately reflect the true predictive performance of the model under this evaluation setting.

**Figure 2.**
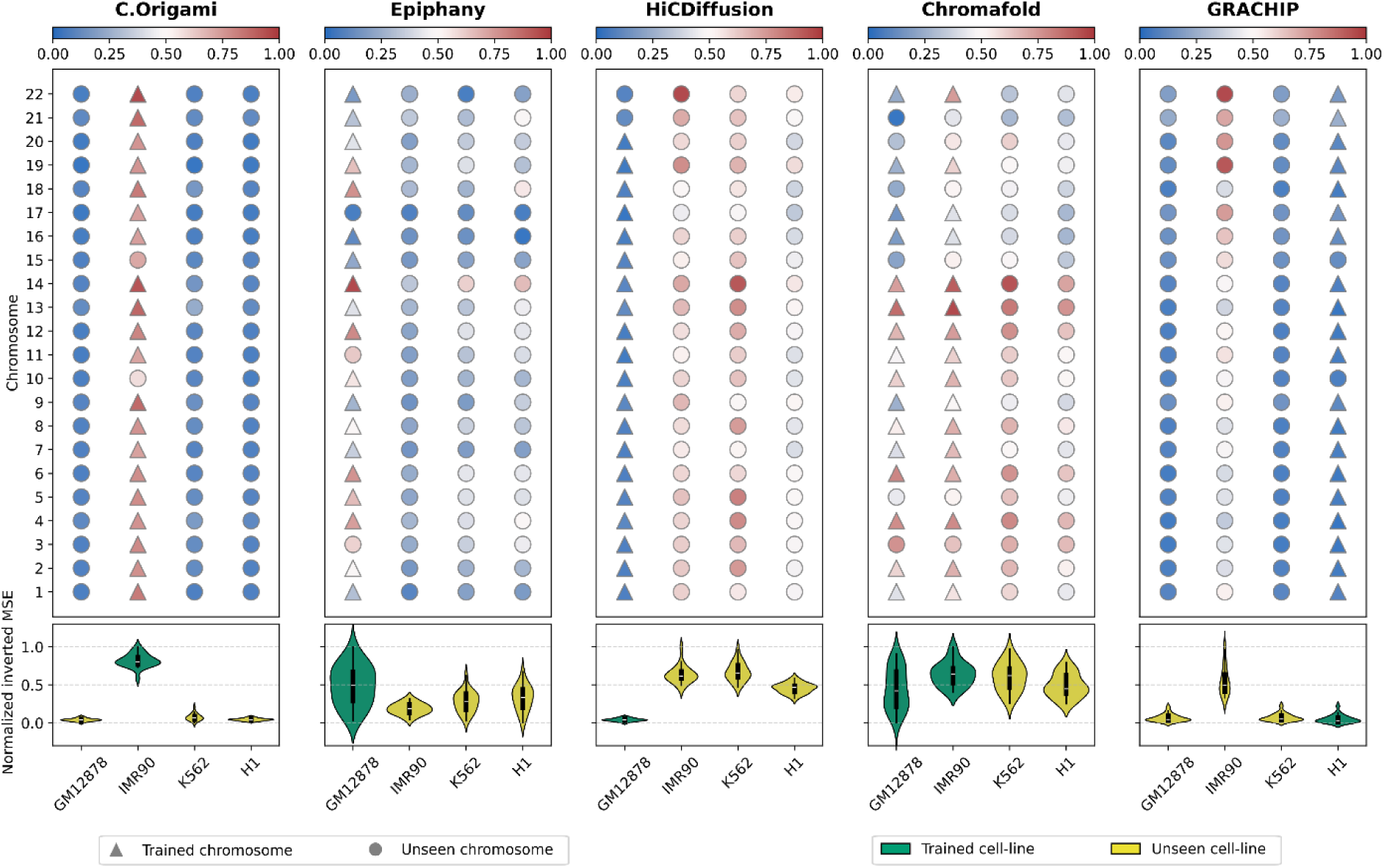
The performance of five Hi-C prediction models using normalized inverted MSE. Inverted MSE is scaled between 0 and 1 with higher values indicating better performance. Top: Bubble plot. The y-axis shows the 22 chromosomes, and the x-axis represents the cell lines. Colors are scaled independently for each cell line. Triangles indicate trained chromosomes, and circles indicate unseen chromosomes. Warmer (red) colors represent higher performance, while cooler (blue) colors indicate lower performance. Bottom: Violin plot showing the distribution of model performance within each cell line. The y-axis denotes the performance scores, and the x-axis corresponds to the cell lines. Green indicates trained cell lines, and yellow indicates unseen cell lines.

Comparing the models based on the insulation score correlation, the results were more consistent and easier to interpret (Figure 3). First, the variance across chromosomes and cell lines appeared more stable and followed a tighter distribution, rather than spanning the extremely wide range observed with MSE. Second, the relationship between training and unseen data aligned with expectations, with performance on the training cell line being slightly higher than performance on unseen cell lines. Third, although C.Origami showed reduced generalization to unseen cell lines, its insulation score correlation remained within a reasonable range (approximately 0.5), suggesting decreased performance rather than the apparent complete failure implied by the normalized MSE results. A similar trend was also observed for the O/E Hi-C correlation (Supplementary Figure S1)

**Figure 3.**
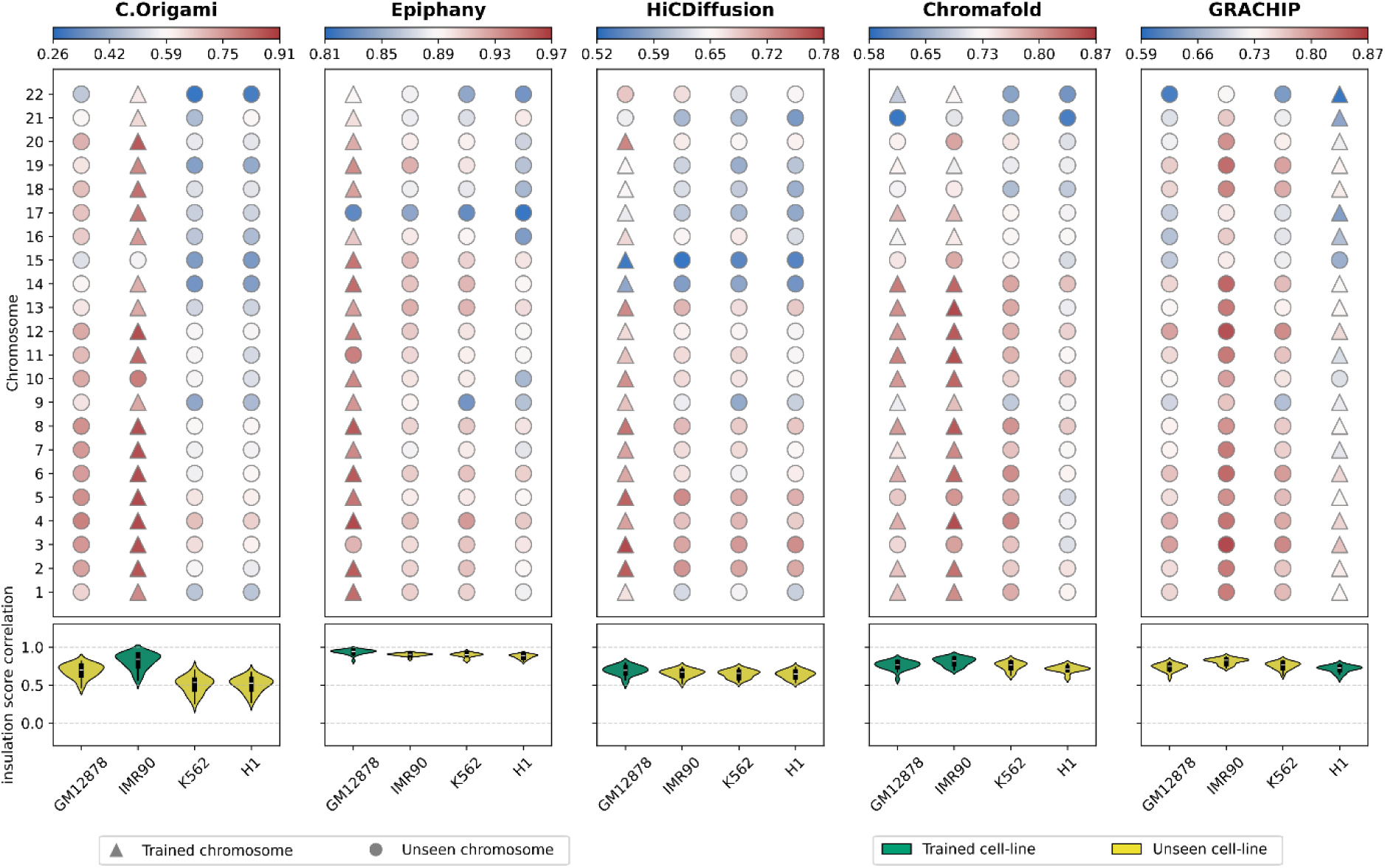
The performance of 5 Hi-C prediction models using insulation score correlation. Top: Bubble plot. The y-axis shows the 22 chromosomes, and the x-axis represents the cell lines. Colors are scaled independently for each cell line. Triangles indicate trained chromosomes, and circles indicate unseen chromosomes. Warmer (red) colors represent higher performance, while cooler (blue) colors indicate lower performance. Bottom: Violin plot showing the distribution of model performance within each cell line. The y-axis denotes the performance scores, and the x-axis corresponds to the cell lines. Green indicates trained cell lines, and yellow indicates unseen cell lines.

This discrepancy indicated that MSE was not well-suited for comparing these models and is not appropriate as a universal comparison metric because the models are trained with different optimization objectives. Epiphany directly minimizes a pixel-wise mean squared error (MSE) between predicted and experimental Hi-C maps (optionally combined with an adversarial loss). In contrast, C.Origami is trained to reproduce structural features of chromatin organization, with performance evaluated using measures such as insulation score and distance-stratified correlation rather than strict intensity matching. ChromaFold is optimized to predict normalized interaction signals and significant contacts from accessibility patterns, while GRACHIP learns interaction relationships through a graph-based representation of chromatin interactions. HiCDiffusion further departs from reconstruction-based training by employing a generative diffusion framework that prioritizes realistic map structure instead of minimizing pixel-wise error. Because these models minimize different objectives during training, an external MSE metric often rewards models trained for numerical reconstruction and penalizes those trained to recover structural or biological features. Therefore, MSE cannot fairly represent real predictive performance across models. Moreover, MSE quantified only pixel-level differences between contact matrices and therefore did not directly measure whether biologically meaningful structures (such as TAD boundaries) were preserved. Finally, because MSE is unbounded (ranging from 0 to infinity), min–max normalization and score inversion could distort interpretation; a low inverted normalized score reflected poorer relative performance within the benchmark rather than true model failure. For these reasons, subsequent evaluations focused primarily on insulation score correlation and observed-over-expected (O/E) Hi-C correlation, which have been widely used to assess the structural similarity and biological validity of predicted Hi-C contact maps in previous studies. In particular, prior work evaluating models such as C.Origami, Epiphany, and related methods relied on insulation profiles, normalized Hi-C similarity, and downstream interaction detection rather than pixel-wise reconstruction error.

Another key observation was that model generalization varied substantially (Figure 3). C.Origami performed best on its training cell line (IMR90), achieving insulation and O/E correlations above 0.9, but its accuracy dropped sharply for unseen lines such as K562 and H1. In contrast, Epiphany maintained high performance across all cell lines, with both correlation metrics exceeding 0.9, demonstrating strong transferability of learned chromatin features. Despite relying solely on DNA sequence, HiCDiffusion also generalized well, producing stable correlations above 0.8 across datasets. ChromaFold showed consistent and robust generalization, maintaining correlations above 0.85 even for unseen cell lines. Although trained on GM12878 and IMR90, its performance on IMR90 and H1 slightly exceeded that on GM12878, suggesting that the model captured transferable regulatory features rather than overfitting. GRACHIP achieved lower overall performance but showed minimal variation across cell lines, indicating stable, though less accurate, generalization.

#### b. Cross-model comparative analysis of Hi-C prediction performance

Subsequently, the models were compared against one another across 22 chromosomes in a specific cell line to identify the strongest performer (Figure 4). The figure reorganizes the previously obtained results and displays the models side by side to enable direct performance comparison. As can be seen, Epiphany achieved the highest overall performance in all metrics, averaging around 0.90 and 0.85, respectively, across all cell lines. ChromaFold followed closely, with strong consistency and minimal variation. C.Origami, again, displayed clear overfitting on the trained cell line, while HiCDiffusion achieved notable accuracy from sequence data alone, although its insulation score correlations were slightly lower than its observed/expected (O/E) Hi-C correlations. GRACHIP ranked lowest, particularly in O/E correlation, around 0.5, reflecting weaker fine-scale reconstruction, though it retained moderate TAD-level (insulations core) accuracy.

**Figure 4.**
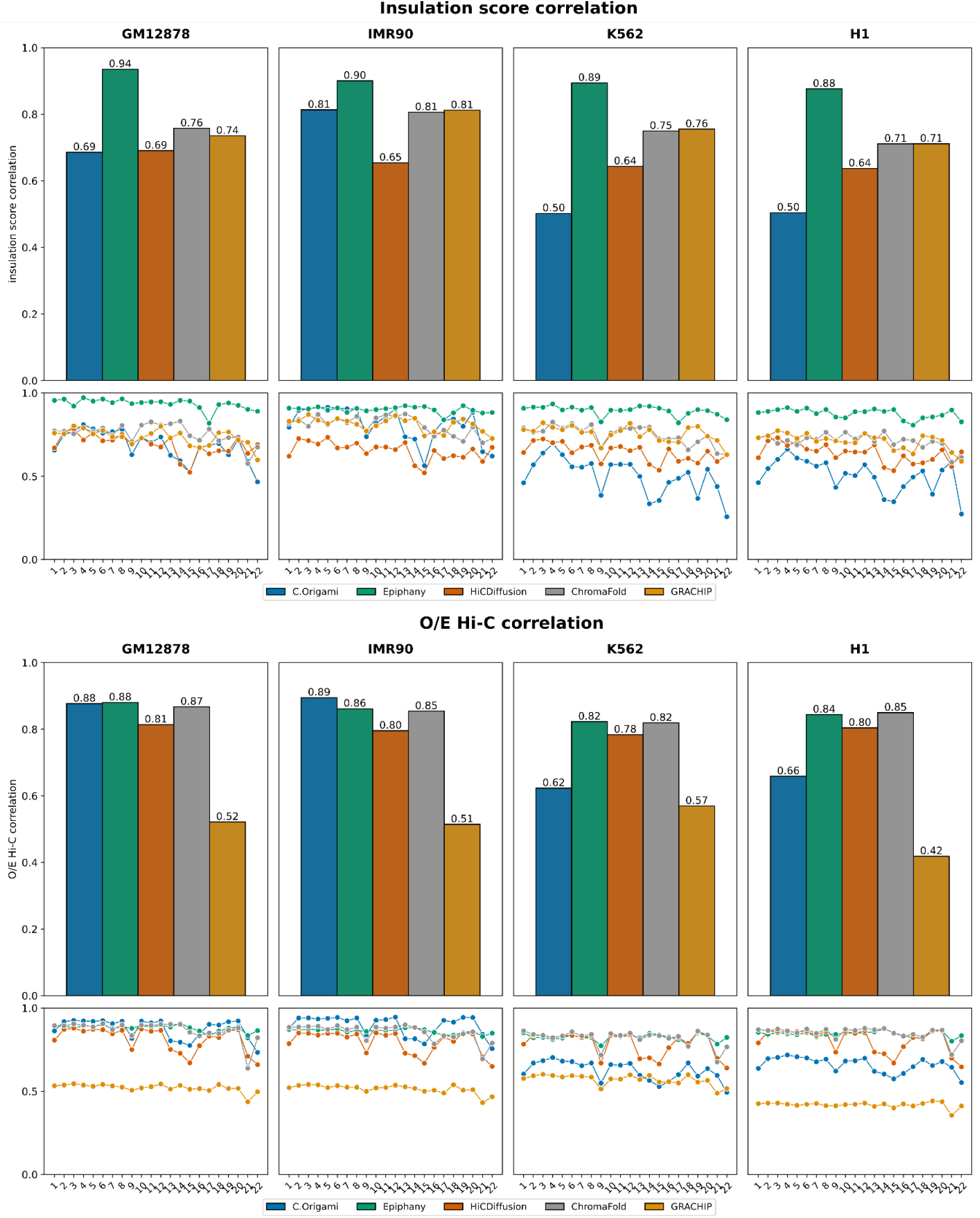
Comparison of model performance for each dataset using insulation score correlation and O/E Hi-C correlation. Top: Bar plot - The y-axis shows the performance score, and the x-axis lists the models, each represented by a distinct colour. Bottom: Line plots - The y-axis shows the performance score across chromosomes, while the x-axis lists chromosome indices. Each line corresponds to a different model, illustrating how model performance varies across genomic regions.

When evaluating performance across individual chromosomes, it was observed that most models exhibited similar trends, consistently struggling with chromosomes 9, 15, 22. This pattern appeared regardless of the specific chromosomes used during training or held out for testing, suggesting that these particular chromosomes might possess structural complexity that were more challenging for models to learn and generalize from. However, despite this challenge, stronger models such as Epiphany and ChromaFold were still able to achieve relatively high scores on these difficult chromosomes.

#### c. Ablation analysis

Ablation testing was performed to quantify the contribution of different input data modalities to Hi-C prediction accuracy, thereby identifying which biological signals were most informative for each model. Specifically, the test was conducted to determine how individual input modalities influence Hi-C prediction accuracy and to investigate whether different regulatory signals provide overlapping information. We systematically removed single modalities or biologically related modality groups and measured the resulting change in prediction performance, thereby assessing potential redundancy and compensatory effects among the input tracks. Due to models with many inputs, only individual modalities and biologically relevant modality groups were tested on four chromosomes (5, 6, 10, and 15). HiCDiffusion was excluded because it relies solely on DNA sequence.

Figure 5 shows the insulation score–based ablation results for Epiphany across input modalities, cell lines, and chromosomes. Corresponding ablation results for the other models and additional evaluation metrics were provided in the Supplementary Material (Supplementary Figure S3 - S13). For clarity, Table 1 summarized the most important input modalities for each model, as determined by ablation results using both insulation score correlation and observed/expected (O/E) Hi-C correlation. Following ablation testing, C.Origami showed a pronounced dependence on CTCF input: removing the CTCF track led to a drastic reduction in performance, indicating that it was the most critical modality for accurate prediction. Notably, the combined use of CTCF binding and ATAC-seq signals further improved predictions, indicating that chromatin accessibility provides complementary information to CTCF binding. In contrast, DNA and ATAC-seq signals alone contributed little in unseen cell lines, indicating potential overfitting to the training data. Similarly, Epiphany’s performance dropped sharply when CTCF binding was removed (insulation correlation <0.8), whereas excluding DNA or histone marks had minimal effect (Figure 5). For ChromaFold, ablation revealed that co-accessibility information was indispensable: removing this feature led to near-zero performance, indicating that the model depends on relationships between accessible regions rather than accessibility peaks alone. This observation is consistent with the original study, which reported that co-accessibility derived from scATAC-seq provides a substantial improvement over pseudobulk accessibility alone. In addition, removal of the CTCF binding signal also reduced performance, suggesting that CTCF binding provides complementary structural information that supports accurate prediction. GRACHIP relied heavily on DNA, with its removal reducing performance to near zero; ATAC-seq also contributed modestly to insulation accuracy. Across models, ablation effects were consistent across chromosomes, suggesting global rather than region-specific dependencies. From all the tests, it revealed that CTCF binding emerged as the most influential feature in most of models, highlighting its essential role in defining chromatin architecture (Table 1).

**Figure 5.**
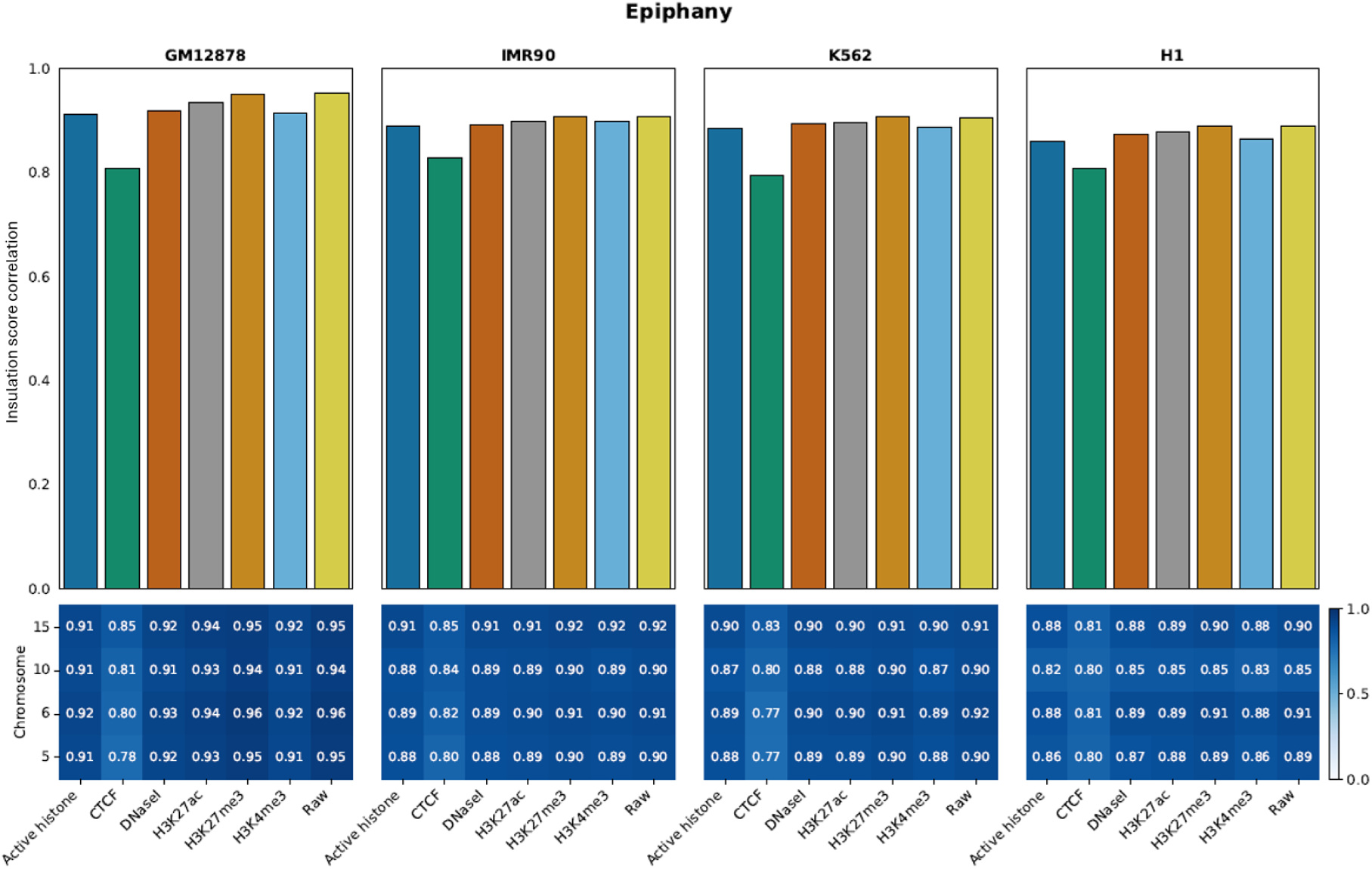
Epiphany ablation performance across modalities, cell lines, and chromosomes, using insulation score correlation. Top: Bar plot showing ablation results; the x-axis lists the removed modality, and the y-axis shows insulation score correlation. Bottom: Heatmap with chromosomes on the y-axis and removed modalities on the x-axis; each cell’s color and value indicate the corresponding performance.

**Table 1.** Summary of important modalities in each model.

| Model | Important modalities |
| --- | --- |
| C.Origami | 1. CTCF ChIP-seq |
|  | 2. bulk ATAC-seq |
| Epiphany | 1. CTCF ChIP-seq |
| ChromaFold | 1. scATAC-seq |
|  | 2. CTCF motif scores |
| GRACHIP | 1. DNA sequence |
|  | 2. bulk ATAC-seq |

### Image quality evaluation

This section evaluated the quality of predicted Hi-C maps using the Fréchet Inception Distance (FID), which measures similarity between real and predicted images by comparing the statistical distributions (mean and covariance) of their feature embeddings (Methods). C.Origami, ChromaFold, and GRACHIP produced blurred or low-contrast maps, reflected in high FID scores exceeding 20. In some cases, trained cell lines even yielded higher FID values than unseen ones, consistent with these models’ optimization for numerical rather than perceptual accuracy.

By contrast, Epiphany and HiCDiffusion achieved significantly better image quality. Epiphany consistently produced the lowest FID scores, which means near-realistic reconstruction of experimental Hi-C maps. HiCDiffusion achieved intermediate performance (FID around 10), due to its generative diffusion process. Overall, the GAN-based Epiphany generated the most visually accurate and biologically plausible contact maps (Figure 6).

**Figure 6.**
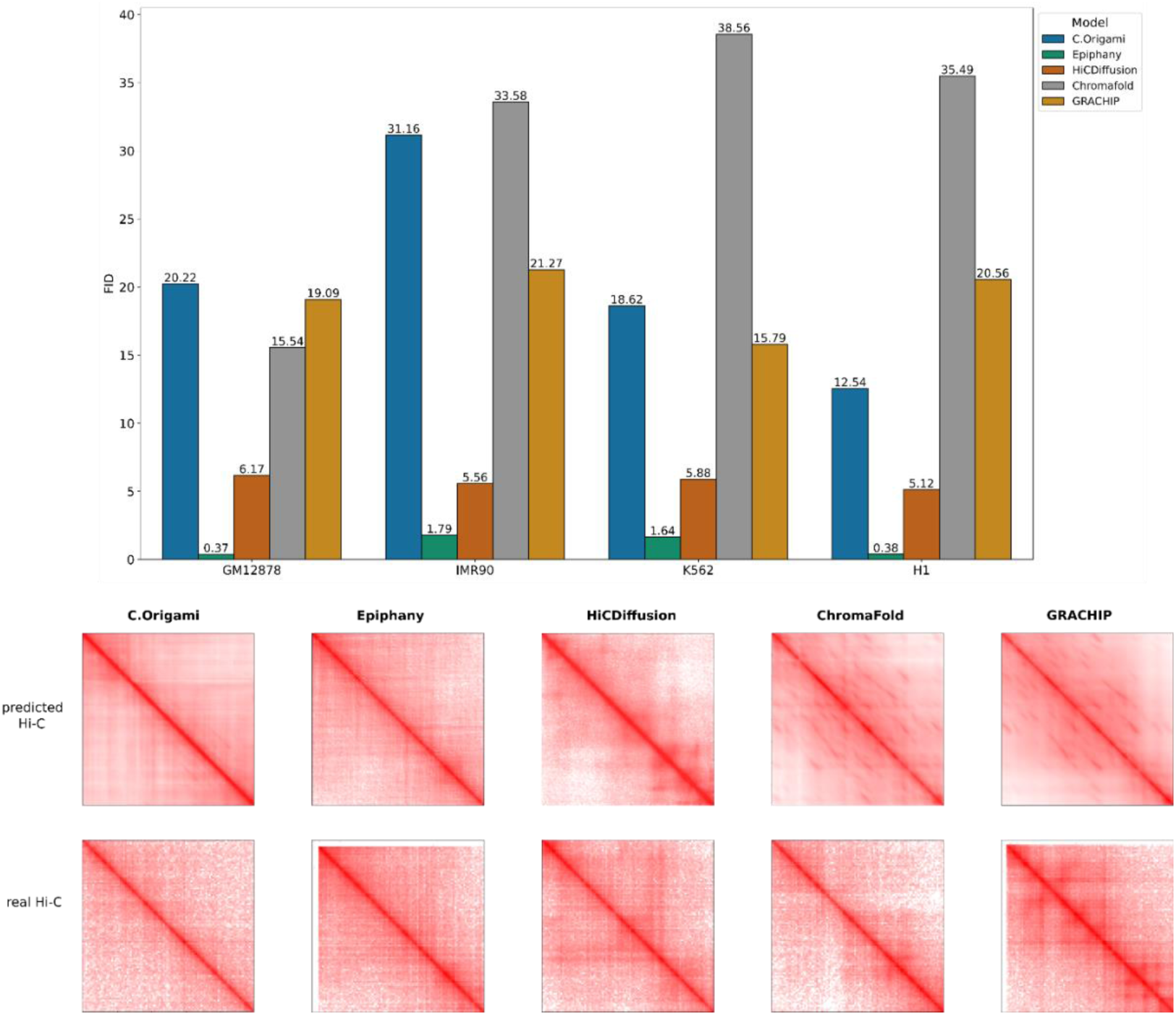
Visual realism of predicted Hi-C contact maps across cell lines. Top: Bar plot showing Fréchet Inception Distance (FID) for each model across cell lines. Bottom: Representative predicted contact maps from each model compared with the corresponding experimental ground-truth Hi-C maps.

### Downstream analysis: loop detection

Chromatin loop detection was used to evaluate the biological relevance of the predicted Hi-C maps. Chromatin loop detection is a downstream Hi-C analysis task that aims to identify specific long-range point-to-point interactions between genomic loci, which appear as focal enrichment “dots” above the local background in Hi-C contact matrices. These loops often reflect structural or regulatory interactions, such as enhancer–promoter contacts mediated by architectural proteins (e.g., CTCF/cohesin), therefore provide a biologically meaningful readout of chromatin organization. To evaluate whether predicted Hi-C maps preserved such biologically relevant features, four loop-calling tools, LASCA [23], HiCCUPS [24], SIP [25], and Mustache [26], were compared in the application to both experimental and predicted maps.

#### a. Functional evaluation of loop detection across models and loop-caller tools

To assess the biological relevance of the detected chromatin loops, loop anchors were compared with independent transcriptional regulatory evidence from three complementary external resources. Curated transcription factor (TF)–target regulatory interactions were obtained from CollecTRI [27], which compiles literature-supported regulatory relationships. Functional downstream targets of TF perturbation were retrieved from KnockTF 2.0 [28], a database of gene expression changes following TF knockdown or knockout experiments. Direct TF–DNA binding evidence was obtained from ChIP-Atlas [29] using ChIP-seq datasets specific to the K562 cell line. These datasets represent distinct levels of regulatory support: CollecTRI provides prior knowledge of regulatory interactions, KnockTF captures functional effects of TF perturbation, and ChIP-Atlas provides cell-type-specific physical TF binding. Rather than claiming direct causality, we evaluated whether predicted chromatin loops were supported by any of these independent regulatory evidences. A loop was considered biologically supported when at least one anchor overlapped a TF binding site (ChIP-Atlas), a TF-regulated gene (KnockTF), or a curated TF–target interaction (CollecTRI). This integration enabled both quantification of loop recovery and assessment of their potential functional relevance. (Methods).

As expected, loops detected from true Hi-C data showed the highest counts and served as the reference set (Table 2). Among predicted contact maps, Epiphany and C-Origami produced the largest number of loops, whereas GRACHIP detected substantially fewer, consistent with its lower structural accuracy. However, loop quantity did not directly correspond to biological support. Loops overlapping K562-specific TF binding sites from ChIP-Atlas were more frequent than those supported only by curated regulatory associations from CollecTRI, indicating that cell-type-specific binding provides stronger support for chromatin looping than general regulatory knowledge. This observation is consistent with established models in which chromatin loops are mediated by active transcription factors in a context-dependent manner. Several models recovered loops enriched for TFs known to be active in K562 cells, including CTCF, RAD21, and NKRF. CTCF and RAD21 are central architectural proteins involved in loop extrusion and maintenance of topologically associating domains, whereas NKRF functions as a transcriptional repressor of immune-related genes [27, 28]. Many predicted loops connected TF binding regions to nearby gene targets, including EIF4G2 and AP2A2 (NKRF-associated), SVIL (RAD21-associated), and SLIT1 and S1PR1 (CTCF-associated), which are relevant to hematopoietic cell function. Nevertheless, loops simultaneously supported by TF binding, regulatory association, and validated downstream targets were rare even in true Hi-C data, reflecting the stringent criteria required to define a functional regulatory chromatin interaction.

**Table 2.**
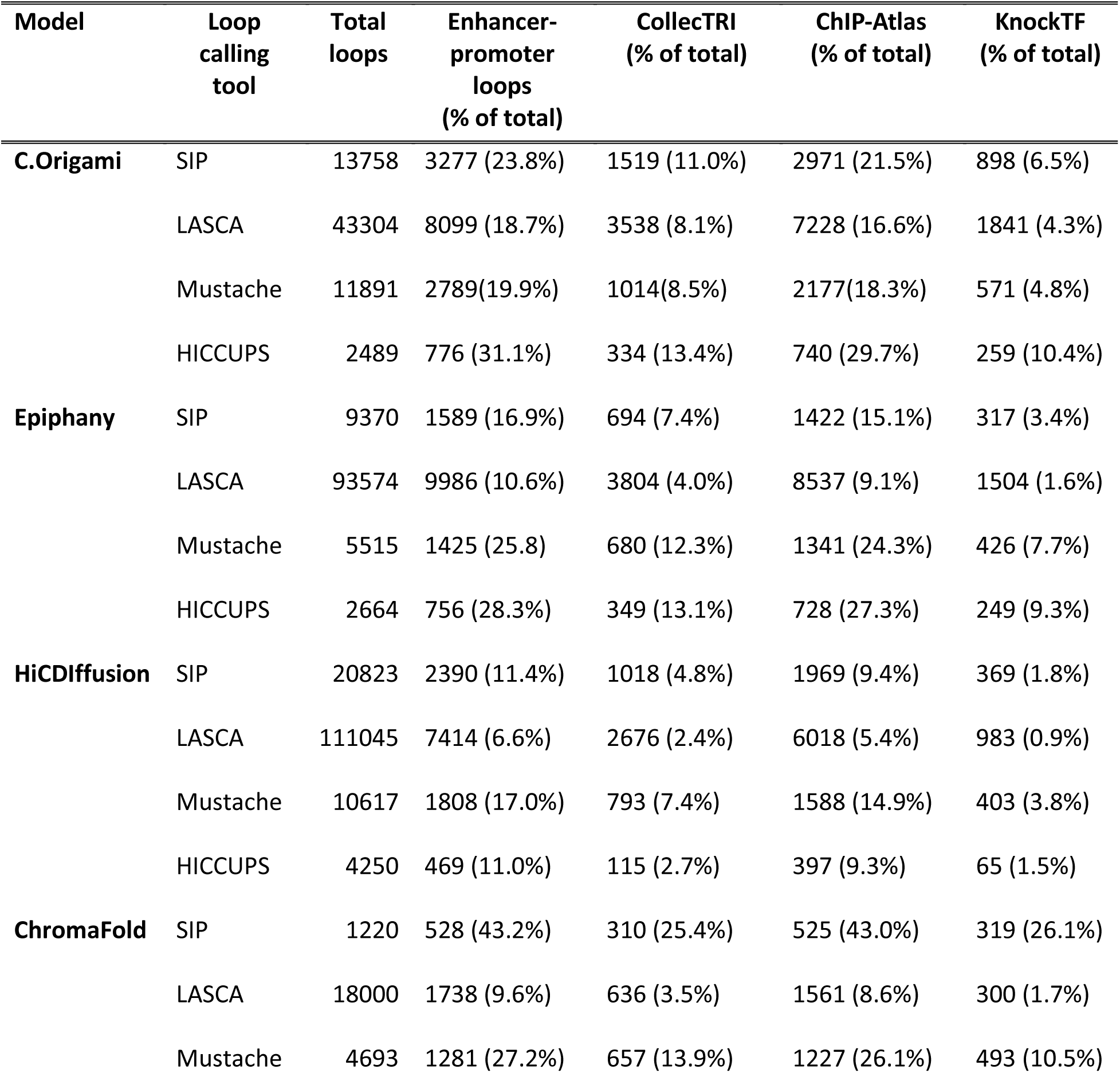
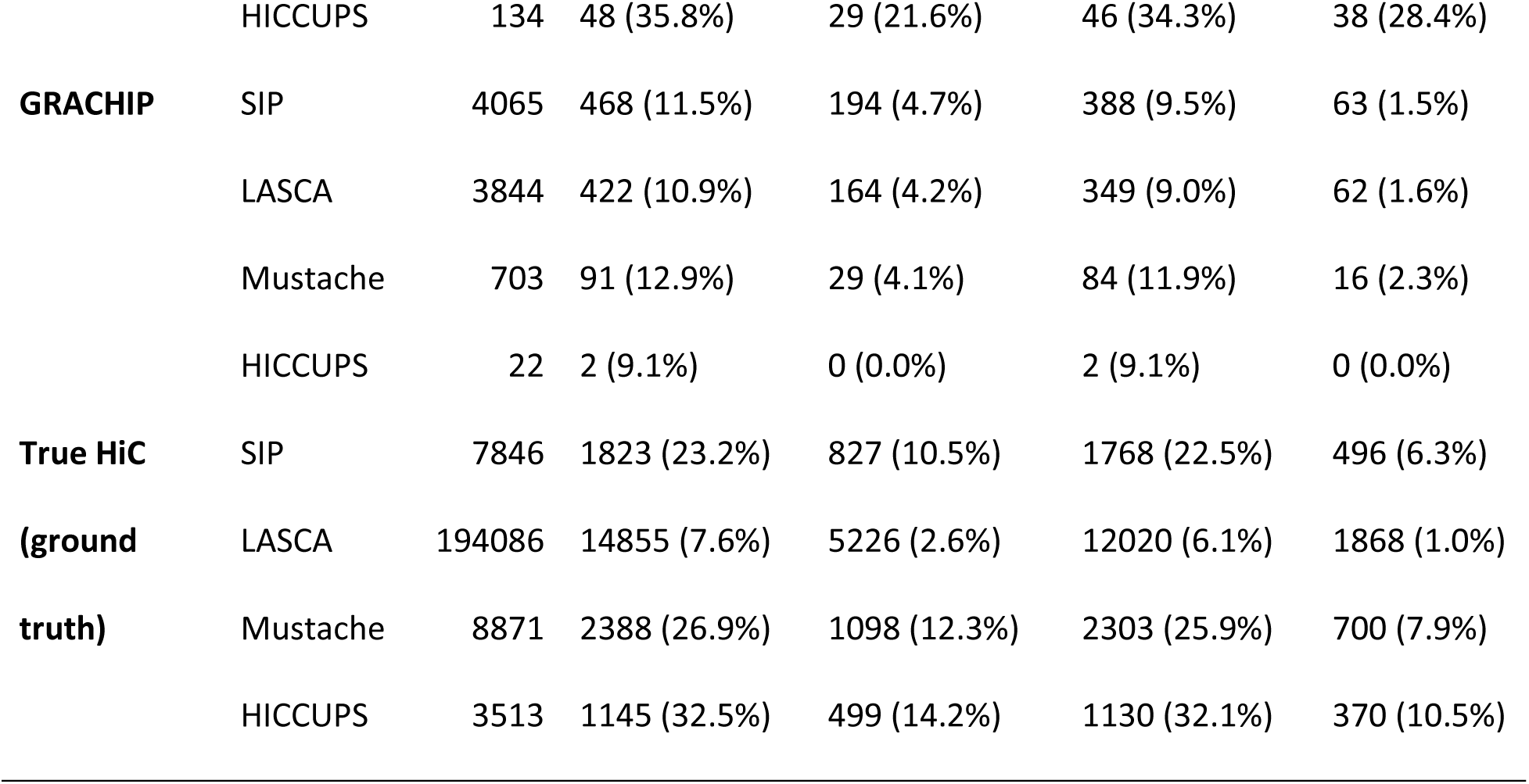
Results of loop detection for each model and specified tools in cell line K562.

Among loop callers, LASCA detected the largest number of loops but exhibited limited overlap with regulatory evidence, suggesting high sensitivity but lower specificity. In contrast, Mustache and SIP detected fewer loops but a higher proportion of biologically supported interactions: approximately 25% overlapped K562-specific TF binding sites and 10% overlapped curated TF regulatory interactions, indicating improved biological precision. HiCCUPS, which relies on sharp focal interaction peaks, performed poorly on GRACHIP maps and detected only 22 loops.

#### b. Cross-model loop overlap using individual loop caller

The biological relevance of predicted Hi-C maps was evaluated by comparing chromatin loops identified from five prediction models against loops derived from the true Hi-C map. For consistency, all loop calls were performed using the same loop-calling tool within each comparison. A loop was initially considered overlapping only if both of its anchors matched those of a loop in another dataset.

C.Origami and Epiphany consistently achieved the highest overlap between predicted biologically supported loops and loops derived from the real Hi-C map across most loop-calling tools (Figure 7). Epiphany’s strong performance was expected given its high-resolution contact maps and high visual fidelity. Interestingly, C.Origami also performed well despite producing comparatively blurred contact maps and demonstrating limited generalization. This suggests that key latent chromatin interaction patterns were still preserved in C.Origami predictions, allowing loop callers, particularly LASCA and Mustache, to recover biologically meaningful interactions even from more diffuse signals. Therefore, visual sharpness alone was not sufficient to fully explain the biological utility of a predicted Hi-C map.

**Figure 7:**
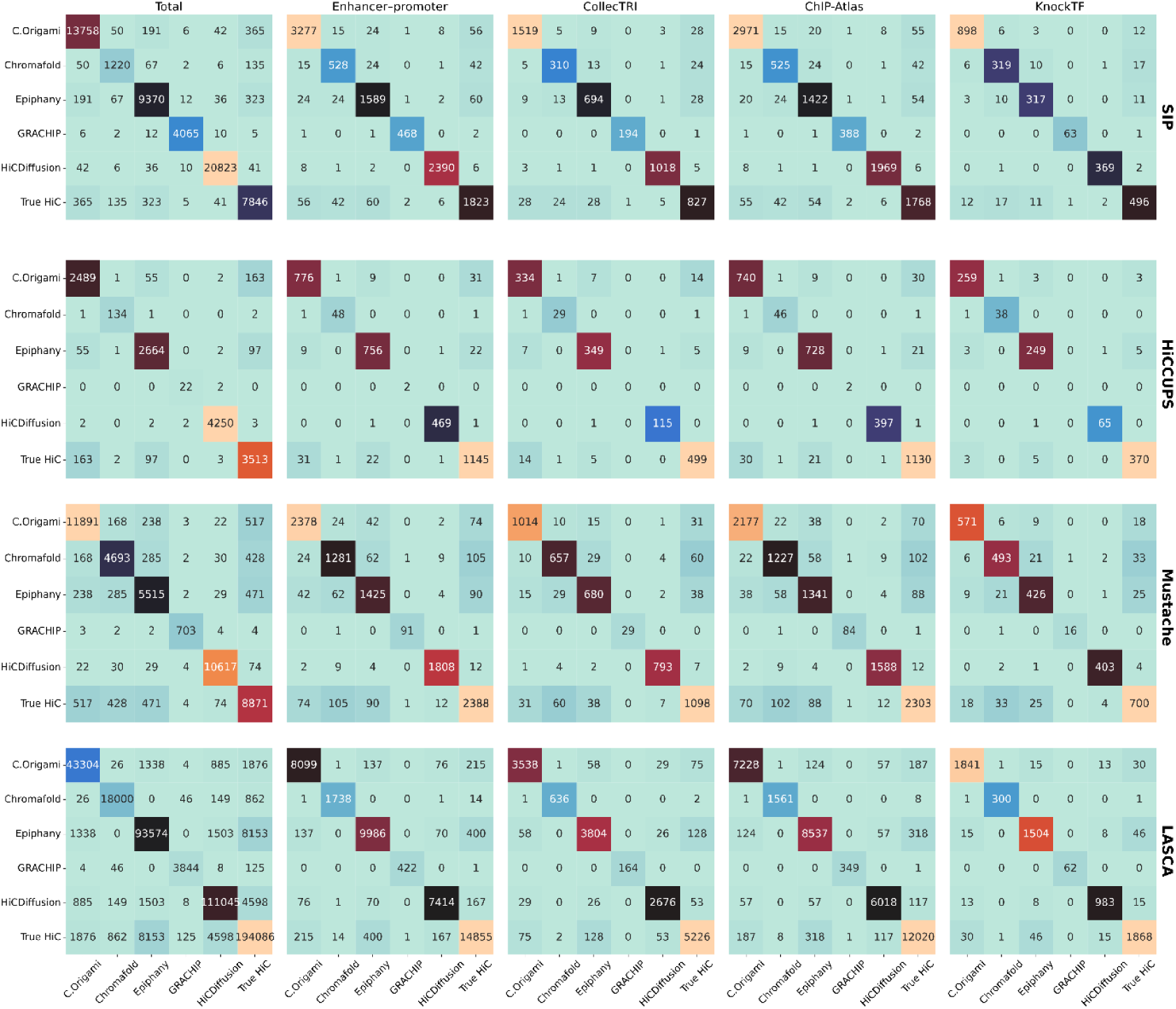
Loop overlap between five models and the true Hi-C map across different loop callers. Heatmaps show the number of overlapping chromatin loops identified between pairs of models and/or ground-truth Hi-C data. In each heatmap, both the x- and y-axes represent the predicted Hi-C maps from different models as well as the true Hi-C dataset. Each cell indicates the number of overlapping loops between the corresponding pair. Color intensity represents the magnitude of overlap, ranging from blue (lowest overlap) to red (highest overlap). Each row corresponds to a different loop-calling algorithm (Mustache, SIP, LASCA, and HiCCUPS). Columns represent different subsets of loops, ordered from left to right: (1) total loops detected by each loop caller (including both biologically meaningful and non-biological loops), (2) loops overlapping enhancer–promoter interactions, (3) loops supported by collectTRI, (4) loops supported by ChIP-Atlas, and (5) loops supported by the KnockTF database.

To quantify loop-calling performance, predicted loops were first compared to loops detected in the true Hi-C map. Loops present in both the predicted and true Hi-C maps were defined as true positives, loops detected only in the predicted map as false positives, and loops present only in the true Hi-C map as false negatives. Using these definitions, precision and recall were calculated for each combination of prediction model and loop-calling tool, and the results were summarized in a two-dimensional precision–recall space (Figure 8). In addition to structural agreement with the true Hi-C loops, we evaluated biological support using three independent regulatory evidence sources: curated TF–target interactions (CollecTRI), functional TF perturbation targets (KnockTF 2.0), and K562-specific TF binding sites (ChIP-Atlas). Across all loop callers, Epiphany and C.Origami consistently ranked among the top-performing models in both precision and recall, indicating robust performance regardless of the loop-calling method used. ChromaFold showed more variable behaviour: it achieved high precision with SIP and obtained the best precision and recall among models when using Mustache, but performance dropped substantially when using other loop callers. Similar trends were observed when evaluating all detected loops and other categories of biologically supported loops (Supplementary Figure S15). Collectively, these results indicate that Epiphany and C.Origami provide the most stable and reproducible performance across multiple loop-calling tools.

**Figure 8.**
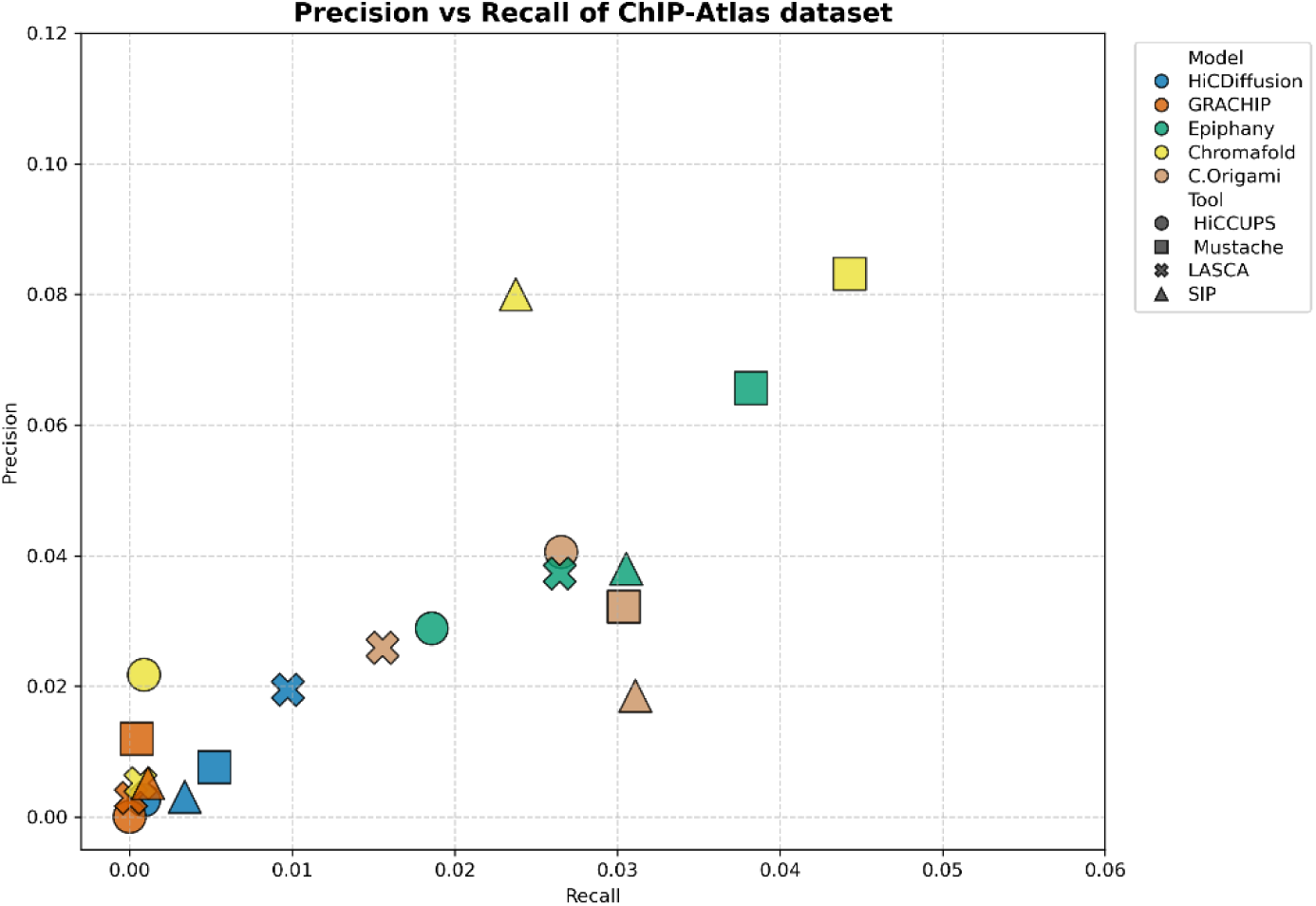
Precision–recall analysis of chromatin loops supported by ChIP-Atlas dataset. The y-axis indicates precision, and the x-axis indicates recall, using loops derived from the true Hi-C map as the ground truth reference. Each marker represents one combination of a Hi-C prediction model and a loop-calling tool. Marker colour denotes the prediction model, while marker shape denotes the loop caller.

Despite these differences, it is important to note that absolute precision and recall values remained low across all models (both metrics typically < 0.15), indicating the inherent difficulty of recovering true chromatin loops from predicted contact maps. To complement structural accuracy, we evaluated the biological support of detected loops. For each loop caller, we calculated the proportion of detected loops whose anchors overlapped independent regulatory evidence from CollecTRI, KnockTF 2.0, or K562-specific ChIP-Atlas binding sites (Figure 9), This metric represents the fraction of predicted loops supported by known or functional regulatory interactions relative to the total number of loops detected, thereby distinguishing potentially functional loops from background structural detections. Using this measure, C.Origami and Epiphany exhibited distributions similar to those of the true Hi-C map across loop callers. This suggests that, although the exact overlap with the ground-truth loop set was limited, these models maintained a comparable ability to generate loops enriched for biological regulatory interactions. In other words, Epiphany and C.Origami produce fewer ground-truth-matching loops, but the loops they do enable are biologically meaningful at a rate comparable to real Hi-C, supporting their utility for downstream biological interpretation.

**Figure 9.**
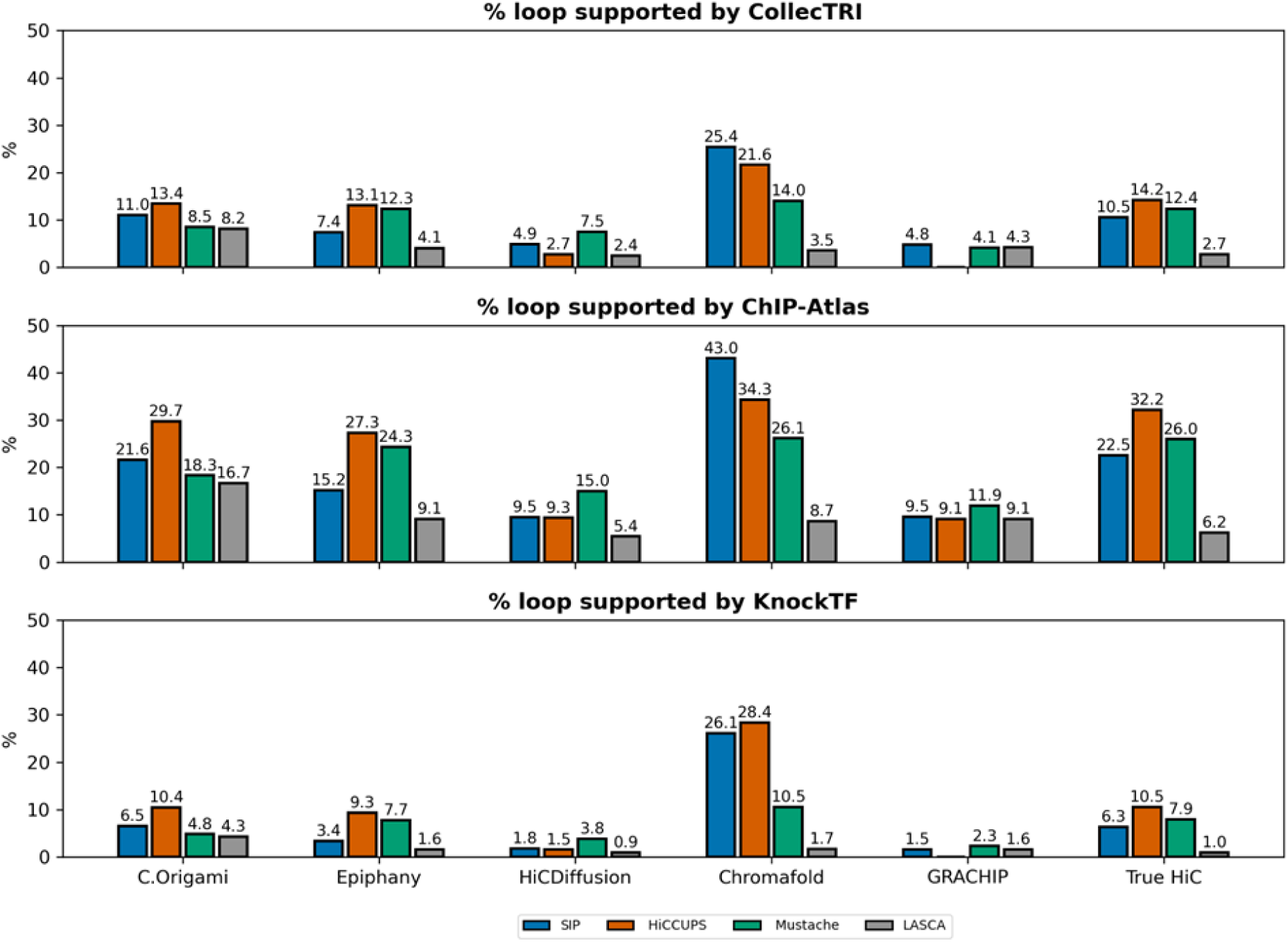
Percentage of biologically supported loops among all detected loops across models and loop callers. The y-axis represents the percentage of biologically supported loops relative to the total number of detected loops. Each panel contains six groups of bars, corresponding to the five prediction models and the true Hi-C map. Bar colour indicates the loop-calling tool used to detect loops.

#### c. Cross-tool loop overlaps within each predicted model

Loop-calling tools also influenced overlap outcomes. HiCCUPS detected fewer overlapping loops due to its dependence on sharp signal peaks, whereas LASCA, Mustache, and SIP were more tolerant to noise and better captured biologically relevant structures. This highlighted the importance of using flexible, noise-resilient tools for analysing predicted Hi-C.

To better understand the behaviour of different loop-calling tools when applied to the same predicted Hi-C map, the overlap of loops identified by SIP, HICCUPS, LASCA, and Mustache within each individual model was evaluated (Supplementary Figure S14). The analysis revealed limited consistency across tools, only a small fraction of loops, particularly enhancer–promoter or TF-associated loops, were shared by all four. Models producing higher-quality predictions, such as Epiphany and C.Origami, exhibited greater cross-tool agreement, indicating that more accurate structural reconstructions yielded more reproducible loop calls. In contrast, models with noisier outputs, like GRACHIP, showed minimal overlap due to inconsistent or spurious detections.

Among the tools, Mustache and SIP demonstrated the highest overlap, likely due to their shared computer vision–based algorithms that emphasize local spatial patterns. LASCA and HiCCUPS, though both clustering-based, differed in behaviour: LASCA detected the most loops but with low biological precision (only around 10% overlapped with known regulatory features), while HiCCUPS remained overly conservative at 10 kb resolution, missing many valid interactions. Biologically meaningful loops identified by three or more tools were rare but often involved enhancer–promoter pairs with TF binding, suggesting that multi-tool consensus loops were highly functionally relevant.

In addition, as shown in Figure 9, HiCCUPS detected the fewest loops among the four tools, yet the fraction of loops overlapping known biological regulatory features was among the highest. This trend was observed not only for the ground-truth Hi-C map but also for most predicted contact maps (except for ChromaFold), with Mustache generally showing the next best performance. This pattern was consistent with previous reports indicating that both HiCCUPS and Mustache performed robustly at 10 kb resolution, where their detection strategies tended to prioritise high-confidence interactions [32]. Interestingly, ChromaFold showed a distinct behaviour: although HiCCUPS and Mustache still yielded high biological precision, SIP consistently achieved the best performance in most cases. Moreover, the fraction of biologically supported loops in ChromaFold was unexpectedly comparable to, or even higher than, that observed in the ground-truth Hi-C map, despite ChromaFold producing fewer loops than true Hi-C and fewer than other high-performing models such as Epiphany and C.Origami. These results suggested that ChromaFold tended to produce a smaller set of loop candidates, yet this set was preferentially enriched for biologically supported interactions and reproducible across loop-calling methods.

## Discussion

### Summary of model characteristics

In this manuscript, five deep learning models, C.Origami, Epiphany, HiCDiffusion, ChromaFold and GRACHIP, were systematically evaluated for their ability to predict Hi-C contact maps and recover biologically meaningful chromatin features, such as loops. These models differed in their architectures, input modalities, and training strategies, resulting in variable performance in prediction accuracy, generalizability, image quality, and biological interpretability. For clarity, each model was compared based on five key aspects: prediction accuracy, generalization across cell lines, input type (single-omic or multi-omics), visual quality of contact maps, and capacity to detect chromatin loops, particularly enhancer–promoter interactions.

C.Origami demonstrated high accuracy in the cell line on which it was trained [21]; however, its performance dropped markedly when applied to unseen cell types, indicating limited generalizability. The model integrates genomic sequence with epigenomic features such as ATAC-seq and CTCF binding. Ablation analysis further revealed a strong dependence on the CTCF track, while other modalities contributed less. Although the visual quality of its predicted maps was relatively low and appeared blurred, the model still captured relevant biological patterns, particularly for loop detection. This observation is consistent with the original study, which identified CTCF as a key determinant of genome organization and loop formation. Accordingly, despite blurred contact maps, C.Origami recovered biologically meaningful chromatin structures, including CTCF–CTCF and promoter–enhancer loops.

Epiphany showed strong and balanced performance and generalized well to new cell types. This agrees with the original study, which specifically designed the model to overcome the limitation of sequence-only predictors that fail to generalize across cell types [18]. This capability is likely driven by its use of multiple epigenomic tracks, including DNase/ATAC-like accessibility, CTCF, and histone modifications such as H3K27ac. Furthermore, by incorporating a GAN-based framework, Epiphany produced visually sharp contact maps that closely resembled experimental Hi-C data. Consequently, Epiphany performed well in detecting both general and biologically supported loops. One limitation, however, is its restricted window size of 1 Mb, which allows detection of interactions up to this distance but may miss longer-range interactions.

HiCDiffusion, which uses only genomic DNA sequence as input, performed surprisingly well [17]. However, sequence-based prediction inherently lacks cell-state information, and prior studies have shown that sequence-only models tend to predict similar structures across cell types because DNA sequence does not encode dynamic regulatory states. The diffusion-based generative framework primarily improves visual quality by reducing blurring in predicted heatmaps and optimizing image realism, as reflected by improvements in the FID. Therefore, although HiCDiffusion produced visually clear maps, its biological specificity should be interpreted cautiously.

ChromaFold achieved moderate accuracy but showed good cross-cell-type generalization. Unlike C.Origami and Epiphany, it relies primarily on single-cell chromatin accessibility and CTCF motif information rather than multiple bulk epigenomic assays [19]. The original study demonstrated that co-accessibility between accessible regions reflects chromatin looping events and enables prediction of cell-type-specific interactions. Previous benchmarks also reported performance comparable to C.Origami and even improved performance when appropriate CTCF information was included. Our ablation results, showing a strong dependence on co-accessibility features, are consistent with this interpretation. The model’s lighter architecture and reduced modality requirements may also reduce overfitting relative to heavily parameterized multi-omics models. Although the predicted maps appeared blurred and the visual quality was limited, ChromaFold was still able to recover chromatin loops, including both general and context-specific interactions.

GRACHIP showed weaker predictive performance in our evaluation despite being designed to improve generalization. The model incorporates DNA sequence, ATAC-seq, and CUT&RUN-derived regulatory features, including CTCF and histone marks, and additionally encodes prior chromatin interaction intensity using a graph neural network [20]. The authors proposed that incorporating interaction relationships would enhance cross-cell-line prediction; however, they also noted that many predictive models, including C.Origami, exhibit performance variability across different cellular contexts. Our results suggest that although the graph-based representation may help capture interactions, it did not translate into improved Hi-C reconstruction accuracy or loop recovery in this benchmark setting.

We summarized the ranking performance of each model across four evaluation categories: accuracy, generalizability, visual quality, and loop detection (Figure 10). Overall, Epiphany ranked highest across all evaluation criteria. C.Origami performed strongly in accuracy and loop detection but showed poor generalization across different cell types. Despite relying only on genomic DNA sequence, HiCDiffusion achieved moderate performance in both generalizability and loop detection. ChromaFold showed consistently moderate performance across all categories, whereas GRACHIP demonstrated the lowest overall performance despite using the largest number of input modalities.

**Figure 10.**
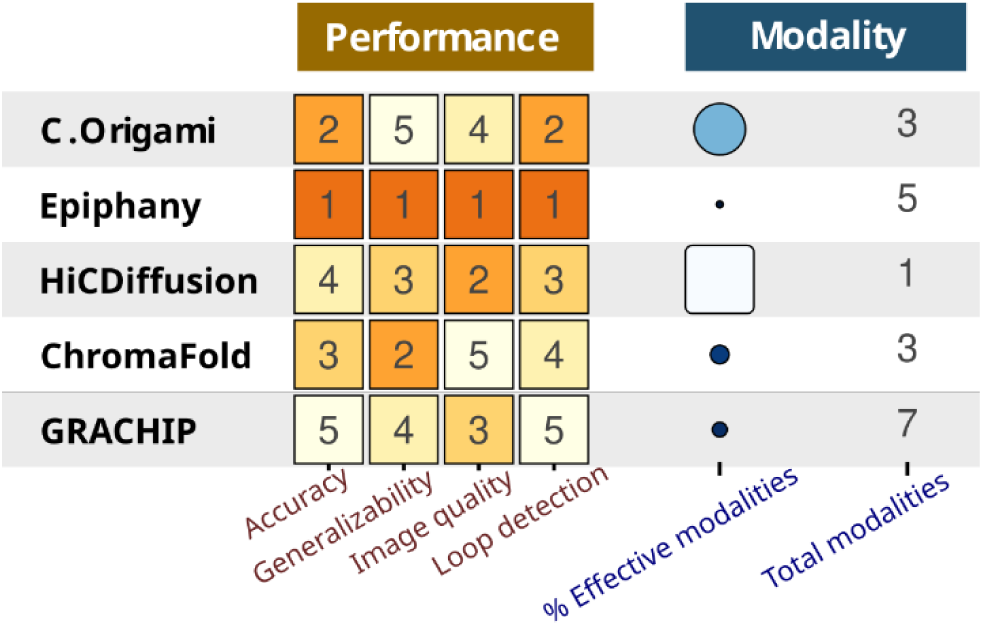
Summary of the evaluation characteristic of each chromatin contact prediction model. Accuracy was ranked based on the average values of two biological metrics, insulation score correlation and O/E Hi-C correlation, calculated across the training cell lines. Generalizability was evaluated using the same metrics averaged across previously unseen cell lines. Visual quality was ranked according to the average FID values across all cell lines. Loop detection performance was determined by the number of biologically meaningful loops predicted by each model relative to the ground truth Hi-C map, averaged across all loop-calling tools. The number of input omics modalities used by each model was also summarized, together with the proportion of modalities contributing measurably to performance, referred to as the “% effective modalities”.

### Insights from the comparative analysis

From the neutral benchmark, several key insights emerged. More input modalities did not necessarily guarantee better performance. Multi-omic models often learned disproportionately from one dominant signal, typically CTCF, while neglecting others , as shown by the ablation analysis in Figure 5 and Supplementary Figure S3-S15, where removal of the CTCF track produced the largest performance decrease compared with other modalities. HiCDiffusion, despite using only genomic sequence, performed comparably or even better than some multi-omic models, demonstrating the importance of model architecture and training strategy over the number of input features.

CTCF binding was a critical input feature across models [33]. As a central player in organizing 3D chromatin architecture, CTCF defines loop anchors and TAD boundaries [31]. Models incorporating experimentally measured CTCF binding generally achieved improved structural reconstruction, particularly for boundary and loop features. However, previous studies also noted that CTCF alone is insufficient to capture cell-type-specific chromatin organization [30]. Additional regulatory context, such as histone modifications or co-accessibility between regulatory elements, was often required to recover accurate and biologically meaningful interaction patterns. Thus, while CTCF provides essential structural information, effective Hi-C prediction depends on combining it with complementary regulatory signals rather than relying on it as a single determinant of genome architecture.

As noted above, many models incorporate multiple input modalities, yet only a subset of these signals substantially influences performance. This suggests that current architectures may not fully exploit the available information and may introduce unnecessary complexity. Importantly, beyond DNA sequence and chromatin accessibility (e.g., ATAC-seq), other epigenomic signals such as histone modifications and transcription factor binding are not as broadly available across cell types, conditions, or species, which may limit the practical applicability of such models. In addition, these epigenomic features are highly context-dependent, raising questions about whether signals derived from one cellular or disease context can be effectively transferred to others. Future work should therefore focus on developing approaches that more effectively integrate complementary regulatory signals while accounting for their availability and context specificity, and on balancing model complexity with informative input features. Improving how modalities are utilized may enhance both prediction robustness and biological interpretability.

MSE proved to be a limited evaluation metric. It did not adequately capture biological relevance or map quality and was highly dependent on model architecture and loss function. For example, models that directly optimized MSE (e.g., C.Origami) often showed artificially low error on training data, while models using adversarial or generative losses (e.g., Epiphany, HiCDiffusion) exhibited higher MSE despite producing biologically realistic and visually accurate maps. Therefore, MSE alone was insufficient to judge model performance. Although MSE is widely used in general machine learning applications, it was not commonly adopted for evaluating Hi-C prediction models. Among the reviewed models, only a few, such as C.Origami, used MSE to assess accuracy, primarily for internal comparisons within the model rather than across different model [21]. In C.Origami’s case, MSE was supplemented with additional evaluation methods, including insulation-based metrics and correlation-based measures, to better reflect biological relevance. In contrast, most comparative studies between models, such as those involving HiCDiffusion vs C.Origami, GRACHIP vs C.Origami, or Epiphany vs Akita, generally avoided MSE altogether. Instead, they focused on metrics more informative for assessing biological and structural fidelity [17, 18, 20]. These typically included correlation-based metrics, such as Pearson and Spearman correlation coefficients, which measured linear and rank-based similarities between predicted and experimental Hi-C matrices, respectively. Other frequently used metrics assessed the ability of models to replicate distance-dependent decay patterns in Hi-C data, such as the O/E ratio and stratum-adjusted correlation coefficient (SCC), which controlled for genomic distance effects [34]. Additionally, metrics based on higher-order chromatin structures, including insulation score correlation and TAD boundary recovery rate, were employed to evaluate the accuracy of TAD predictions [35]. In this study, we employed not only MSE but also two biologically meaningful evaluation strategies: distance-dependent decay analysis using O/E correlation and TAD-level evaluation using insulation score correlation. Further refinement could include metrics like the stratum-adjusted correlation coefficient or distance-specific correlation measures, which offer a more nuanced understanding of how well predicted contact maps reflect the complex spatial organization of the genome.

Another important consideration is that the models evaluated in this study were originally trained on different cell lines. To maintain fidelity to the published versions, we used the original training settings and verified reproducibility through limited retraining. While this ensures that the models were assessed as intended for practical use, it introduces a potential limitation in comparability due to differences in training data. Notably, following the original papers, ChromaFold was trained on two cell lines, whereas the other models were trained on a single cell line. This may have contributed to differences in performance and generalizability; however, the current analysis does not allow us to disentangle the effects of model architecture from those of training data diversity. Future work could address this by retraining all models on standardized datasets to enable a more controlled comparison.

Finally, the performance of the same deep learning model varied across different chromosomes. For example, the prediction accuracy tended to fluctuate across the 22 chromosomes, with some chromosomes showing peaks in performance while others exhibited noticeable decreases. Interestingly, in the same model, this trend was almost consistent across four different cell lines: H1, GM12878, IMR90, and K562, despite their biological differences. Even when using five different models, all trained on different chromosomes or cell lines, a similar performance pattern was observed. Specifically, a drop in prediction accuracy was consistently seen on chromosomes 9, 15, and 22 across all models and cell lines. This consistent decrease suggested that chromosomes 9, 15, and 22 presented unique challenges for generalization and prediction, regardless of the model architecture or training data. There were several potential biological and technical explanations for this observation. First, chromosome size may play a role. Smaller chromosomes, such as chromosome 22, contains fewer genomic loci and thus generated fewer interaction pairs in Hi-C data [36]. This results in sparser contact maps, which were more difficult for models to learn and accurately predict. Second, the structural complexity of certain chromosomes may affect performance. For instance, chromosomes 9 and 15 are known to contain complex repetitive regions and were hotspots for chromosomal translocations [37]. These structural variations lead to mapping ambiguities and reduced the quality of Hi-C signals, making it harder for models to learn consistent patterns. Third, chromatin interaction also varies across chromosomes: gene-rich chromosomes have denser transcriptional activity and more regulatory interactions, whereas gene-poor chromosomes exhibit sparser patterns [8]. If a model is trained on more chromosomes with rich regulatory interactions and then tested on chromosomes with sparse or fundamentally different interaction landscapes, performance can drop due to poor generalization. These differences directly affect Hi-C signal patterns and may influence how easily a model can learn and generalize across chromosomes. In addition, variability in sequencing depth, differences in chromatin organization (such as TADs and compartments), and uneven distribution of epigenomic features like CTCF binding sites and histone modifications might also contribute to performance differences across chromosomes. Taken together, these findings indicate that both biological properties and technical factors influenced model performance across chromosomes in our benchmark, consistent with known variability in Hi-C data and chromatin organization.

### Loop detection from predicted Hi-C map

Loop detection represents an important biologically meaningful downstream application of predicted Hi-C contact maps. While most previous Hi-C prediction studies primarily evaluated downstream performance using TADs, loop-level interactions have received far less attention. Moreover, when loops were assessed, studies typically relied on a single loop-calling method, such as HiC-DC+, to identify significant interactions. Because different loop callers rely on distinct algorithmic principles, the choice of method can substantially influence loop detection results. To our knowledge, few studies have systematically examined how different loop-calling algorithms perform on predicted Hi-C maps, making this analysis an important contribution of the present work.

Our results show that, although the evaluated prediction models operated at a relatively coarse resolution (around 10 kb), many were still able to recover biologically relevant interactions, including enhancer–promoter loops and TF-associated chromatin contacts. Nevertheless, precision and recall values remained low when predicted loops were directly compared with loops derived from the true Hi-C map. This limited performance is expected for several reasons. First, prediction errors in contact intensity and local structural patterns can introduce noise, resulting in inconsistent loop detection across callers. Second, the 10 kb resolution itself imposes a fundamental constraint: at this scale, fine chromatin features, particularly short-range or tightly localized interactions, may be blurred, merged, or lost entirely. In addition, resolution mismatch and boundary uncertainty complicate the mapping of loop anchors between predicted and true Hi-C, further reducing apparent overlap even for biologically related interactions. This indicates resolution as a key limitation for accurate loop recovery. Future work should explore higher-resolution predictions (e.g., 5 kb or 1 kb) or post hoc enhancement methods to improve loop detection.

Another consideration is the choice of the loop-calling tool. Predicted maps often contained noise, blur, or artifacts that are not present in real Hi-C data. Some tools, like HICCUPS, which are sensitive to sharp peaks and clean signals, performed poorly on predicted maps. In contrast, tools based on computer vision (e.g., Mustache, SIP) showed more robust performance, likely due to their tolerance to noise and blurred patterns [32]. Similar observations have been reported in recent loop-detection studies, where algorithm performance depends strongly on signal quality and map resolution[32]. These observations suggest that loop-caller performance depends not only on algorithm design but also on structural pattern of the predicted contact maps.

Importantly, the structural quality of the predicted contact maps appeared to be more critical than the specific choice of loop-calling tool. When a predicted contact map closely resembled the ground truth Hi-C map in both signal characteristics and visual structure, as was observed with models like Epiphany and HiCDiffusion, most loop callers, including sensitive ones like HICCUPS, were able to effectively detect chromatin loops. This indicates that, under high-quality input conditions, different loop-calling algorithms tend to converge on similar biologically meaningful interactions.

It is often assumed that visual fidelity correlates with structural quality; however, an important exception was observed for C.Origami. Although C.Origami was not explicitly optimized to improve the visual realism of predicted contact maps, it still performed strongly in loop detection. Specifically, its overall loop-calling performance was consistently second only to Epiphany and exceeded that of HiCDiffusion, while also producing a substantial number of biologically supported loops overlapping those derived from true Hi-C. Notably, C.Origami enabled recovery of a relatively large number of loops consistently across multiple loop callers, including conservative tools such as HiCCUPS, indicating that its predicted contact maps retained sufficient structural pattern for robust loop identification. These results indicate that loop callers may be robust to certain forms of visual blurring, as long as key underlying chromatin interaction patterns are preserved.

This insight also raises an important question: to what extent does visual fidelity contribute to the overall quality of predicted Hi-C maps for reliable loop detection? In this study, it was difficult to disentangle the effects of visual quality and prediction accuracy, as ,for example, Epiphany performed strongly in both aspects: achieving strong accuracy, generating visually realistic maps, and showing greater overlap with true biological loops. As Epiphany combined accurate prediction with GAN-based refinement that improved map sharpness, the independent contribution of visual quality to loop-calling outcomes could not be determined. Future work could address this by comparing model variants with and without visual refinement under identical loop-calling pipelines to assess whether improved image quality alone enhances biological discovery.

## Conclusion

In conclusion, we systematically benchmarked five deep learning models for Hi-C chromatin contact prediction across four cell lines, evaluating their predictive accuracy, visual fidelity, and ability to support downstream loop detection. Ablation analysis revealed that epigenomic features, particularly CTCF binding and co-accessibility, are key contributors to accurate Hi-C prediction, while many additional modalities provide limited benefit. Among all models, Epiphany performed consistently well across all criteria, demonstrating high accuracy, generalizability, visual quality, and effective loop detection. Furthermore, our results highlight that predicted Hi-C maps can effectively support loop detection.

## Methods

### Data and cell line selection

This study used publicly available datasets from four well-characterized human cell lines: K562, IMR-90, GM12878, and H1-hESC, which were chosen for their extensive publicly available multi-omics data and obtained from the ENCODE and 4DN databases (Supplementary Table S1). IMR-90, GM12878, and H1-hESC were included in the training data of at least one evaluated model, whereas K562 was not. This allowed assessment of model performance on both seen and unseen cell types, providing insight into their generalization ability. Furthermore, the study only used 22 chromosomes for each cell line, excluding sex chromosomes.

The input data used for Hi-C contact map prediction included genomic DNA sequence and various epigenomic data. All genomic sequences were aligned to the hg38 human reference genome. Epigenomic data comprised chromatin accessibility (ATAC or DNase), CTCF binding, and histone modifications such as H3K4me3, H3K27ac, H3K4me1, and H3K27me3 (all by ChIP-seq).

### Deep learning models for comparison

This study used five advanced deep learning models for the prediction of Hi-C contact maps: C.Origami, Epiphany, HiCDiffusion, ChromaFold, and GRACHIP. These models collectively represent a broad range of contemporary deep learning approaches, including CNNs, transformers, recurrent neural networks (RNNs), graph convolutional networks (GCNs), as well as generative techniques such as GANs and diffusion-based models (Supplementary Table S3).

All models were executed using the authors’ publicly available implementations. The software was installed in a Python (3.11) environment with PyTorch as the primary deep learning framework, along with the dependencies specified in each repository. Training was performed on the Ghent University high-performance computing (UGent HPC) cluster using GPU acceleration (NVIDIA GPUs), while inference and downstream analyses were carried out on CPU nodes. Model parameters and preprocessing steps followed the recommendations of the original publications unless otherwise stated.

**C.Origami** [21] predicts Hi-C contact maps by integrating genomic and epigenomic information (hg38), using three inputs: DNA sequence, CTCF binding, and ATAC-seq chromatin accessibility. The genomic sequence was provided in FASTA format and one-hot encoded into five channels (A, T, C, G, and N), which remained constant across all cell types. The ATAC-seq and CTCF data were pre-processed with the Seq-N-Slide pipeline (https://igordot.github.io/sns/) to align raw FASTQ reads, remove low-quality sequences, and produce normalized bigWig signal tracks. These signals were log-transformed to reduce noise and ensure numerical stability before being used as input. The Hi-C training data were processed at 10 kb resolution and downsampled to 8,192 bp to match the model’s output format, with a reversible natural logarithm transformation applied to stabilize training. The model architecture consisted of two main components: feature extraction and contact map prediction. The feature extraction stage included two one-dimensional convolutional encoders that captured local genomic patterns, followed by a transformer module that modelled long-range dependencies. The transformer, adapted from the BERT architecture, contained eight attention layers with 256 hidden units and eight attention heads, using relative positional embeddings to encode genomic distance. Each input covered a 2.1 Mb genomic region divided into 256 bins and contained seven channels: five from DNA sequence and two from ATAC-seq and CTCF signals. After feature extraction, the transformer output was pairwise concatenated to generate a 256 × 256 × 256 tensor, which was then decoded through five dilated residual blocks to produce a 256 × 256 contact matrix representing predicted chromatin interaction intensities at 8,192 bp resolution. The model was trained using MSE loss on the IMR-90 cell line, with all chromosomes except 10 and 15 used for training; Chromosome 10 served as validation and Chromosome 15 as the test set.

**Epiphany** [18] predicted Hi-C contact maps from epigenomic data representing 3D genome organization, using DNase I hypersensitivity, CTCF binding, and three histone marks (H3K4me3, H3K27ac, and H3K27me3). All input data were processed as described in the original method paper through a unified pipeline in which replicate BAM files for each modality were merged and converted into bigWig signal tracks using the bamCoverage tool with a 10 bp bin size and RPGC normalization to ensure consistent signal intensity across samples. The Hi-C training data were processed at 10 kb resolution and normalized using Hi-C-DC+, which applied a negative binomial regression model to compute observed-over-expected interaction ratios while accounting for genomic distance, GC content, and restriction fragment size. These normalized interaction values served as the model’s training targets. Epiphany employed a sliding window approach in which each 1 Mb stripe of the contact matrix was predicted using a 1.4 Mb window of input signals, including 200 kb of padding on both sides. During training, the model predicted 200 consecutive stripes per forward pass, corresponding to an input tensor of size 5 channels × 3.4 Mb × 200 stripes, with a 10 kb stride between windows. The model architecture was based on a GAN consisting of a generator and discriminator. The generator contained multiple convolutional layers for local feature extraction followed by a bidirectional LSTM layer to capture long-range dependencies across stripes, producing realistic contact map predictions through a final fully connected layer. The discriminator, composed of convolutional layers, enforced an adversarial objective to distinguish real from generated maps. The combination of MSE and GAN losses enhanced prediction accuracy and structural realism. Epiphany was trained on the GM12878 cell line, with chromosomes 3, 11, and 17 held out for validation and testing to assess generalization.

**HiCDiffusion** [17], unlike multimodal models that integrated epigenomic data, relied solely on sequence features derived from the hg38 human reference genome, excluding telomeric and centromeric regions to avoid repetitive and noisy sequences. The model took a 2,097,152 base-pair (2 Mb) genomic window as input, one-hot encoded into five channels representing A, T, C, G, and N bases. Each input corresponded to a predicted Hi-C contact matrix of 256 × 256 bins at 8,192 bp resolution. The architecture followed a two-stage transfer learning framework composed of a pretrained encoder-decoder backbone and a diffusion-based refinement process. The encoder compressed the 2 Mb sequence into 256 bins with 256 feature channels using convolutional layers, residual connections, batch normalization, and ReLU activations, followed by a transformer module with eight attention heads to capture long-range dependencies. The decoder employed dilated convolutions to reconstruct an initial 256 × 256 single-channel contact map, which served as the base prediction. This pretrained encoder–decoder was then integrated into a diffusion refinement stage, where a U-Net denoising network iteratively learned to remove Gaussian noise added to the predicted maps. During training, the model minimized reconstruction error by progressively denoising corrupted contact maps, and during inference, it started from random noise and refined it through learned iterative denoising steps guided by the encoder–decoder features. This approach enabled the generation of visually realistic and structurally coherent contact maps. HiCDiffusion was trained on the GM12878 cell line and used all autosomal chromosomes except Chromosome 22 for testing and Chromosome 21 for validation.

**ChromaFold** [19] used single-cell chromatin accessibility data with CTCF motif scores to predict HiC contact map data. The model leveraged two major features derived from scATAC-seq data: pseudobulk chromatin accessibility and co-accessibility profiles. Pseudobulk accessibility was generated by aggregating single-cell accessibility signals across a population of cells to provide a robust representation of overall chromatin openness. To address the sparsity of scATAC-seq data, metacells, clusters of similar cells, were constructed, and co-accessibility between genomic regions was quantified using the Jaccard similarity of binarized accessibility profiles across metacells, enabling the model to capture interaction patterns at a population level. The CTCF motif score track was computed by scanning genomic DNA sequences with CTCF position weight matrices to identify potential binding sites that contribute to chromatin looping and domain organization. The Hi-C training data were processed at 10 kb resolution and normalized into Z-scores using HiC-DC+, with values clipped between –16 and +16 to reduce the influence of outliers. ChromaFold’s architecture consisted of two parallel feature extractors: the first processed pseudobulk accessibility and CTCF motif score tracks through 15 convolutional layers to capture local context, while the second extracted interaction-specific features from co-accessibility data. Outputs from both extractors were concatenated and passed through a linear prediction layer that estimated contact strengths between a central genomic bin and its surrounding bins within a 2 Mb window, forming a V-shaped stripe prediction. The model was trained in two stages for improved convergence: first optimizing the primary extractor and temporary prediction head, then freezing the learned weights while training the co-accessibility branch and final layer. ChromaFold used 4.01 Mb input windows and was trained on IMR-90 and GM12878 cell lines, with chromosomes 3 and 15 reserved for validation and chromosomes 5, 18, 20, and 21 held out for testing.

**GRACHIP** [20], a graph-based deep learning model for predicting Hi-C contact maps, integrated 7 input modalities: genomic DNA sequences, ATAC-seq data, and five CUT&RUN-derived epigenomic features: CTCF, H3K4me3, H3K4me1, H3K27ac, and H3K27me3, based on the hg38 human reference genome. CUT&RUN data were processed through the 4DN pipeline, while ATAC-seq data followed the ENCODE ATAC-seq pipeline. Both pipelines generated normalized bigWig signal tracks as final outputs. For CUT&RUN, raw reads underwent quality control, adapter trimming, alignment with Bowtie2, and filtering to retain only high-quality, unique reads before producing bigWig tracks. Similarly, ATAC-seq reads were aligned, duplicates and mitochondrial reads were removed, and chromatin accessibility signals were output in bigWig format. DNA sequence data were processed using a pretrained DNABert model, which transformed nucleotide sequences into 768-dimensional embeddings that captured rich contextual information. Hi-C training data were processed with the 4DN Hi-C pipeline at 10 kb resolution, including alignment with BWA, filtering with pair tools, duplicate removal, and normalization of final contact matrices after merging biological replicates. GRACHIP’s architecture combined three modules to model multi-scale genomic dependencies: a three-layer GCN to encode genomic features and structural relationships, a 12-layer Transformer with 12 attention heads to capture long-range dependencies, and a six-layer CNN to extract local interaction patterns. The model produced a 200 × 200 contact matrix representing predicted chromatin interactions within a 2 Mb genomic window at 10 kb resolution. By integrating graph-based representations with sequence and epigenomic features, GRACHIP effectively modelled both linear and spatial genomic contexts. The model was trained on the H1 human embryonic stem cell line, using all chromosomes for training except chromosomes 10 and 15, which were reserved for evaluation.

Most models were unable to predict interactions across the entire length of a chromosome due to memory and computational constraints. Instead, predictions were made on fixed-size windows, commonly 2 Mb in length. Supplementary Table S2, S3, S4 summarize key information for each model, including the input modalities, resolution, cell lines used for training, the specific deep learning techniques employed in each architecture, and the source code of each model.

### Overview of workflow

The workflow began with data collection and preprocessing. Each evaluated model required specific input modalities and preprocessing procedures. To ensure fair and reproducible evaluation, all models were retrained using the original preprocessing steps, input modalities, and training settings described in their original publications. The same cell line and chromosome splits were also maintained for training, ensuring that the retrained models produced results consistent with those originally reported. Following preprocessing, the prepared data were input into each model to generate predicted Hi-C contact maps across five human cell lines. During inference, the same preprocessing protocols were applied to maintain methodological consistency. The predicted contact maps were then compared with experimental Hi-C data from the corresponding cell lines. To assess prediction accuracy, the predicted contact maps were quantitatively compared against the ground truth Hi-C maps of the same cell lines. For image quality assessment, the FID was used to evaluate the visual similarity between predicted and true contact maps. Finally, to evaluate the utility of the predicted maps in downstream tasks, both predicted and ground truth contact matrices were input into loop-calling tools. The resulting loops were compared to evaluate how well each model recovered biologically relevant chromatin interactions. While some original model studies examined structural features such as loops or TADs, they typically relied on a single analysis strategy and did not systematically compare performance across multiple loop-calling algorithms. Here, we extended these approaches by applying several loop callers and directly comparing loop recovery, providing a more standardized assessment of downstream biological utility.

### Evaluation criteria for model comparison

#### a. Accuracy performance

Model performance was evaluated by comparing predicted Hi-C matrices with experimentally derived contact maps using three quantitative measures: MSE, insulation score correlation, and O/E Pearson correlation. MSE quantifies the average squared difference between predicted and ground-truth contact frequencies. Insulation score correlation evaluates whether the model captures local chromatin domain organization (e.g., TAD boundaries) by computing insulation scores along the genome from both predicted and experimental matrices and comparing the resulting insulation profiles using Pearson correlation. Specifically, for each genomic bin, insulation scores are computed within a fixed window (≈500 kb) by summarizing average contacts in upstream, downstream, and central regions, and calculating the ratio 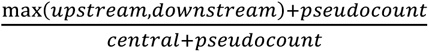. Finally, O/E Pearson correlation measures global agreement after normalizing contact frequencies to account for distance-dependent decay, by flattening the observed/expected-normalized matrices and computing Pearson correlation. Before analysis, all matrices were log-transformed using 𝑙𝑜𝑔_10_(𝑐𝑜𝑢𝑛𝑡 + 1) to normalize data distributions. Comparisons were conducted separately for each chromosome and cell line to assess model robustness across biological contexts. Results were also compared between training and unseen cell lines to evaluate generalization. All performance metrics were summarized under identical conditions to enable direct model comparison.

#### b. Ablation testing

To assess the contribution of each input modality, we performed ablation testing by removing selected features and measuring the resulting change in performance. Unlike Epiphany, which retrained models after masking input tracks, we conducted post-training ablation: model parameters were fixed and removed inputs were replaced with neutral signals. This approach evaluates the model’s dependence on each modality during inference rather than its ability to relearn without it, and also avoids the substantial computational cost and inconsistent training procedures across models. For sequence inputs, nucleotides were replaced with the ambiguous base “N.” For epigenomic features (e.g., ATAC-seq or histone marks), signal values were set to zero while preserving genomic coordinates. Models with few input types (C.Origami and ChromaFold) were tested under all modality combinations, whereas models with more inputs were evaluated using single and grouped ablations. Modalities were grouped by biological function: (1) chromatin accessibility (ATAC-seq, DNase I), (2) transcription factor binding (CTCF), (3) activating histone marks (H3K4me3, H3K27ac, H3K4me1), and (4) repressive histone marks (H3K27me3). HiCDiffusion was excluded because it relies only on DNA sequence. Ablation analyses were performed on chromosomes 5, 6, 10, and 15 as representative genomic regions.

#### c. Image quality

Quantitative metrics alone cannot fully assess the perceptual realism of predicted Hi-C maps. Therefore, FID was used to evaluate the visual quality of predicted Hi-C contact maps. FID is a standard metric for assessing generative models, including GANs and diffusion models, by comparing the distribution of generated images to that of real images in the feature space of a pretrained InceptionV3 network. Specifically, it fits multivariate Gaussian distributions to the extracted feature activations and computes the Fréchet distance between them based on their mean and covariance. Lower FID values indicate greater visual similarity to the experimental Hi-C maps.

#### d. Downstream loop detection analysis

To evaluate the biological utility of predicted Hi-C maps, downstream analysis focused on identifying chromatin loops associated with transcriptional regulation. Four loop-calling tools were used: LASCA [23], HiCCUPS [24], SIP [25], and Mustache [26]. These four loop callers were selected for several reasons. First, they are well-established and user-friendly tools that can be efficiently executed in high-performance computing (HPC) environments. Second, they represent two major categories of loop detection algorithms with two tools selected from each category. Within each group, the selected tools are known to perform optimally at different resolutions, allowing us to evaluate loop detection performance across algorithmic strategies and resolution sensitivities. Among these, LASCA and HiCCUPS were clustering-based methods, while SIP and Mustache applied computer vision techniques. Clustering-based tools used algorithms such as DBSCAN to detect loops by grouping nearby points based on spatial proximity and cluster size, with HiCCUPS identifying sharp focal peaks using a donut-shaped local background, and LASCA employing graph-based community detection to handle complex and sparse data. In contrast, computer vision methods analysed the Hi-C contact maps as images: Mustache detects loops using scale-space theory to identify peaks at multiple scales, and SIP integrates advanced image processing to identify loops and domains by recognizing spatial patterns. The reason for choosing these four tools was to evaluate how predicted Hi-C maps perform in loop detection using two different types of algorithms: clustering-based and computer vision-based methods. Within each category, we selected two tools to capture variation in loop detection sensitivity at different resolutions. HiCCUPS and Mustache were known to predict more loops at 10 kb resolution, while LASCA and SIP tended to detect more loops at higher resolutions like 5 kb [32]. Since the predicted Hi-C maps in this study have a 10 kb resolution, comparing these tools allows assessment of loop detection performance across different algorithms and resolution sensitivities.

For this analysis, loop calling is performed on the predicted and true Hi-C maps (converted into the required “.hic” format using juicer tool with KR normalization). The K562 cell line was chosen, as it was not included in model training, providing an unbiased test of generalization.

After loop calling, detected loops were annotated with enhancer and promoter regions. Enhancers were obtained from EnhancerAtlas 2.0 [38], and promoters were defined as ±2 kb around transcription start sites using GENCODE (GRCh38) annotations. To assess biological relevance, predicted loops were compared with three reference resources: CollecTRI [27], a curated database of transcriptional regulatory interactions; ChIP-Atlas [29], which provides transcription factor binding profiles; and KnockTF 2.0 [28], which contains gene expression data following transcription factor knockdown or knockout. For the KnockTF dataset, differential expression analysis between TF knockdown and control samples was performed using the *limma* package, executed in Python via the rpy2 interface. Only statistically significant TF–target gene pairs (FDR < 0.05) were retained, and these pairs were mapped onto predicted chromatin loops to identify regulatory interactions supported by experimental evidence.

Finally, several comparisons were made:

1. Total number of predicted loops compared to the ground truth loops from the reference Hi-C map.
2. Overlap between enhancer-associated loops predicted by each model and true Hi-C loops.
3. Overlap between predicted loops and curated TF–target regulatory interactions (CollecTRI).
4. Overlap between predicted loops and transcription factor binding evidence (ChIP-Atlas).
5. Overlap between predicted loops and expression-supported TF–target relationships derived from KnockTF 2.0.

Results from all loop-calling tools were then compared to evaluate the performance of predicted Hi-C maps across different loop detection methods. This framework assesses whether predicted chromatin contacts are not only structurally plausible but also supported by independent regulatory and gene expression evidence.

## Declarations

Ethics Approval and Consent to Participate - N/A

Consent of Publication - N/A

Funding - N/A

## Availability of Data and Materials

All datasets used in this study are publicly available. Hi-C datasets were obtained from the 4DN Data Portal [39]. Details of all Hi-C and other omics datasets used in this study are provided in Supplementary Table S1. The code used in this manuscript is available at https://github.com/CBIGR/bulk_hic_benchmark.

## Author Contributions

Contributions according to the CRediT system (https://www.ucl.ac.uk/library/research-support/open-access/credit-taxonomy), HTTN = Hoang Thien Thu Nguyen, VV = Vanessa Vermeirssen): Conceptualization: VV, Data curation: HTTN, Formal analysis: HTTN, Funding acquisition: VV, Investigation: HTTN, VV, Methodology: HTTN, VV, Project administration: VV, Software: HTTN, Supervision: VV, Validation: HTTN, Visualization: HTTN, Writing-original draft: HTTN, VV, Writing-review and editing HTTN, VV. All authors read and approved the final manuscript.

## Acknowledgements

This work was facilitated through the high-performance computing (HPC) infrastructure provided by Ghent University. We thank Jens Uwe Loers for his guidance on HPC computing. We also thank the CBIGR team members for their valuable feedback and suggestions that helped improve the study.

## Additional File

Supplementary document includes additional figures and tables detailing data sources, analysis results, and extended visualizations.

