## Supplementary document for "Evaluation of deep learning tools for chromatin contact prediction"

**Supplementary tables**

**Supplementary Table S1.** Datasets used in this study.

| Modality | Cell line | Database | ID |
|---|---|---|---|
| **HiC** | GM12878 | 4DN | 4DNFIXP4QG5B |
| | IMR90 | 4DN | 4DNFIJTOIGOI |
| | H1 | 4DN | 4DNFIQYQWPF5 |
| | K562 | 4DN | 4DNFI18UHVRO |
| **ATAC** | GM12878 | ENCODE | ENCFF603BJO |
| | IMR90 | ENCODE | ENCFF282RNO |
| | H1 | 4DN | 4DNFICPNO4M5 |
| | K562 | ENCODE | ENCFF077FBI |
| **CTCF** | GM12878 | 4DN | 4DNES6GVE8XZ |
| | IMR90 | ENCODE | ENCFF105FHL |
| | H1 | 4DN | 4DNES1RQBHPK |
| | K562 | 4DN | 4DNESMSEE62M |
| **H3K4me3** | GM12878 | 4DN | 4DNES9CEMVIB |
| | IMR90 | ENCODE | ENCFF376ZIM |
| | H1 | 4DN | 4DNESPE6J9FU |
| | K562 | 4DN | 4DNESV634K6K |
| **H3K27ac** | GM12878 | 4DN | 4DNESOOUQAN7 |
| | IMR90 | ENCODE | ENCFF699OAR |
| | H1 | 4DN | 4DNESIMWCLF8 |
| | K562 | 4DN | 4DNFIEYAZI72 |
| **H3K27me3** | GM12878 | 4DN | 4DNES7W1FL5I |
| | IMR90 | ENCODE | ENCFF366BVS |

| | | | |
|---|---|---|---|
| | H1 | 4DN | 4DNES8TY5P5P |
| | K562 | 4DN | 4DNESTTCK612 |
| | GM12878 | 4DN | 4DNESLYCNMZP |
| **H3K4me1** | IMR90 | ENCODE | ENCFF221MJG |
| | H1 | 4DN | 4DNESRFWR5SV |
| | K562 | 4DN | 4DNES6VYFIZW |
| | GM12878 | ENCODE | ENCSR000EMT |
| **Dnase I** | IMR90 | ENCODE | ENCSR477RTP |
| | H1 | ENCODE | ENCSR000EMU |
| | K562 | ENCODE | ENCSR000EOT |

**Supplementary Table S2.** Summary of training and evaluation of cell lines used for the different models.

| Model | GM12878 | IMR90 | H1 | K562 |
|---|---|---|---|---|
| **C.Origami** | Unseen | Trained | Unseen | Unseen |
| **Epiphany** | Trained | Unseen | Unseen | Unseen |
| **HiCDiffusion** | Trained | Unseen | Unseen | Unseen |
| **ChromaFold** | Trained | Trained | Unseen | Unseen |
| **GRACHIP** | Unseen | Unseen | Trained | Unseen |

**Supplementary Table S3**. Summary of models with modality, resolution, interaction distance and architecture.

| Model | Modality | Resolution | Interaction distance | Architecture |
|---|---|---|---|---|
| **C.Origami** | ATAC,<br>CTCF,<br>DNA | 8,192 bp | 2,097,152bp | CNN,<br>Transformer |
| **Epiphany** | DNase I,<br>CTCF,<br>H3K4me3,<br>H3K27ac,<br>H3K27me3 | 10,000bp | 1Mbp | GAN<br>CNN,<br>Bi-LSTM |
| **HiCDiffusion** | DNA | 8,192 bp | 2,097,152bp | Diffusion,<br>CNN,<br>Transformer |
| **ChromaFold** | CTCF motif scores,<br>Pseudobulk chromatin accessibility,<br>Co-accessibility profiles | 10,000bp | 2Mbp | CNN |
| **GRACHIP** | DNA,<br>ATAC,<br>CTCF,<br>H3K4me3,<br>H3K4me1,<br>H3K27ac,<br>H3K27me3 | 10,000bp | 2Mbp | GCN,<br>CNN<br>Transformer |

**Supplementary Table S4**. Source code link of each chromatin interaction prediction model.

| Model | Source code |
|---|---|
| **C.Origami** | https://github.com/tanjimin/C.Origami.git |
| **Epiphany** | https://github.com/arnavmdas/epiphany.git |
| **HiCDiffusion** | https://github.com/SFGLab/HiCDiffusion.git |
| **ChromaFold** | https://github.com/viannegao/ChromaFold.git |
| **GRACHIP** | https://github.com/Ruoyun-W/GRACHIP.git |

### Supplementary figures

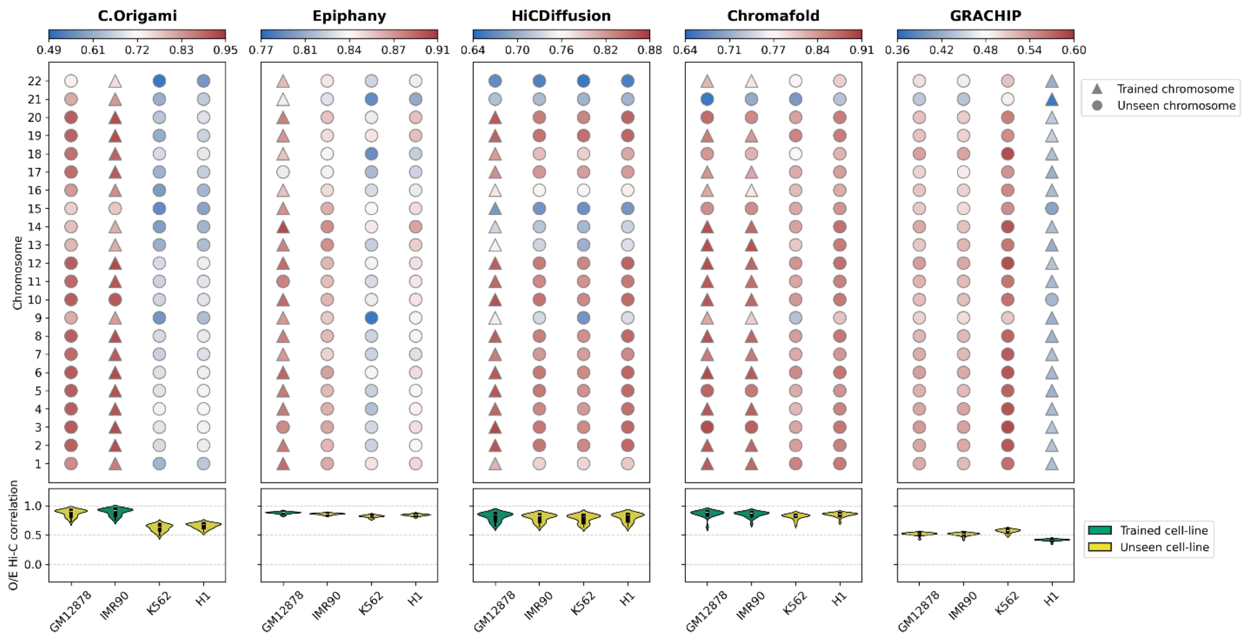


**Supplementary Figure S1.** The performance of 5 Hi-C prediction models using O/E Hi-C correlation. Top: Bubble plot. The y-axis shows the 22 chromosomes, and the x-axis represents the cell lines. Colors are scaled independently for each cell line. Triangles indicate trained chromosomes, and circles indicate unseen chromosomes. Warmer (red) colors represent higher performance, while cooler (blue) colors indicate lower performance. Bottom: Violin plot showing the distribution of model performance within each cell line. The y-axis denotes the performance scores, and the x-axis corresponds to the cell lines. Green indicates trained cell lines, and yellow indicates unseen cell lines.

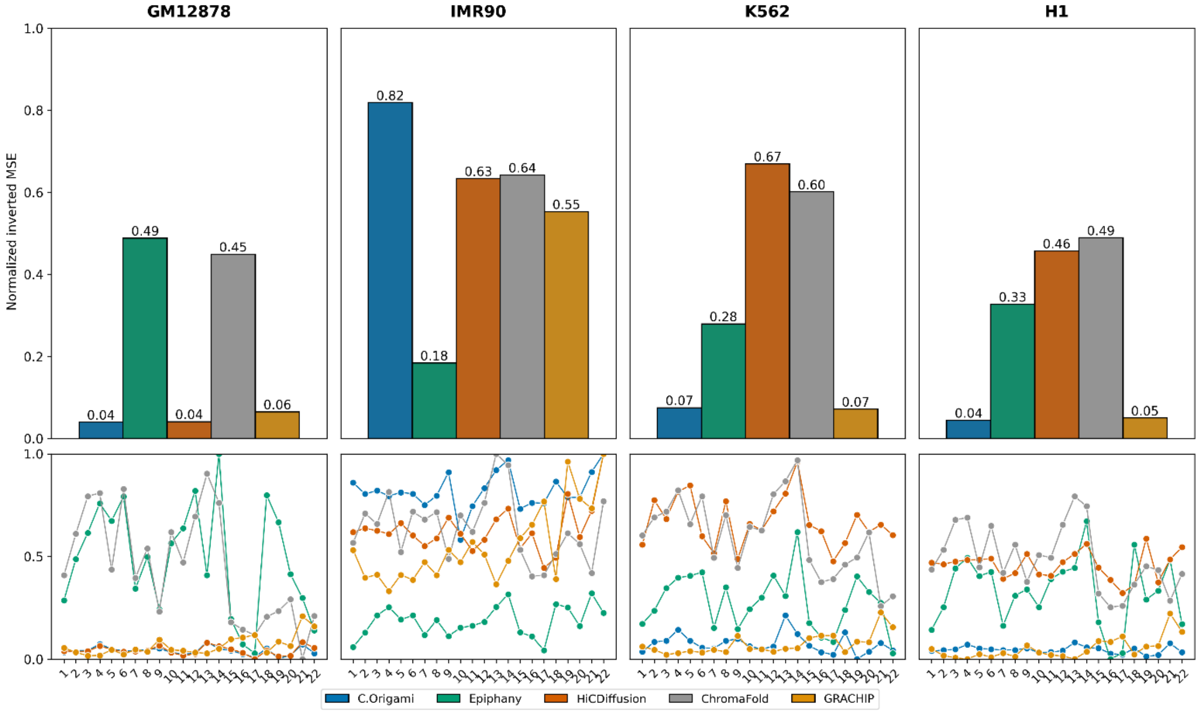


**Supplementary Figure S2.** Comparison of model performance for each dataset using normalized inverted MSE. Top: Bar plot - The y-axis shows the performance score, and the x-axis lists the models, each represented by a distinct colour. Bottom: Line plots - The y-axis shows the performance score across chromosomes, while the x-axis lists chromosome indices. Each line corresponds to a different model, illustrating how model performance varies across genomic regions.

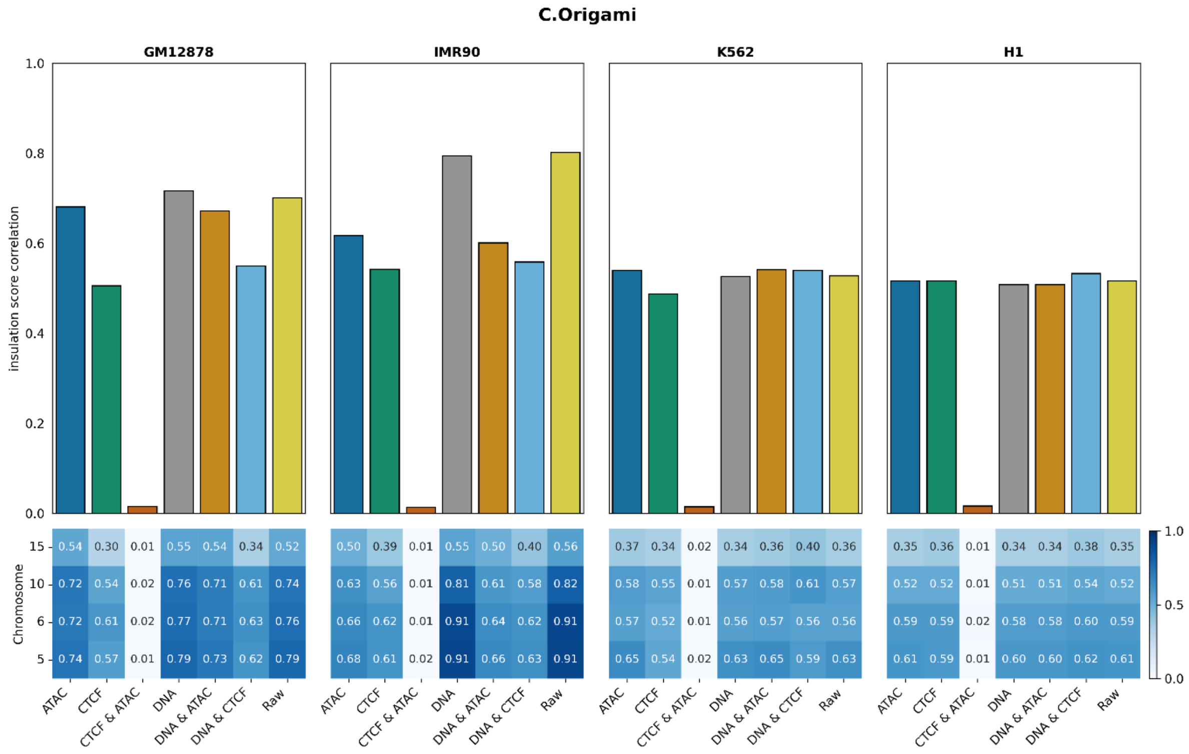


**Supplementary Figure S3.** C.Origami ablation performance across modalities, cell lines, and chromosomes using insulation score correlation. Top: Bar plot showing ablation results; the x-axis lists the removed modality, and the y-axis shows insulation score correlation. Bottom: Heatmap with chromosomes on the y-axis and removed modalities on the x-axis; each cell's color and value indicate the corresponding performance.

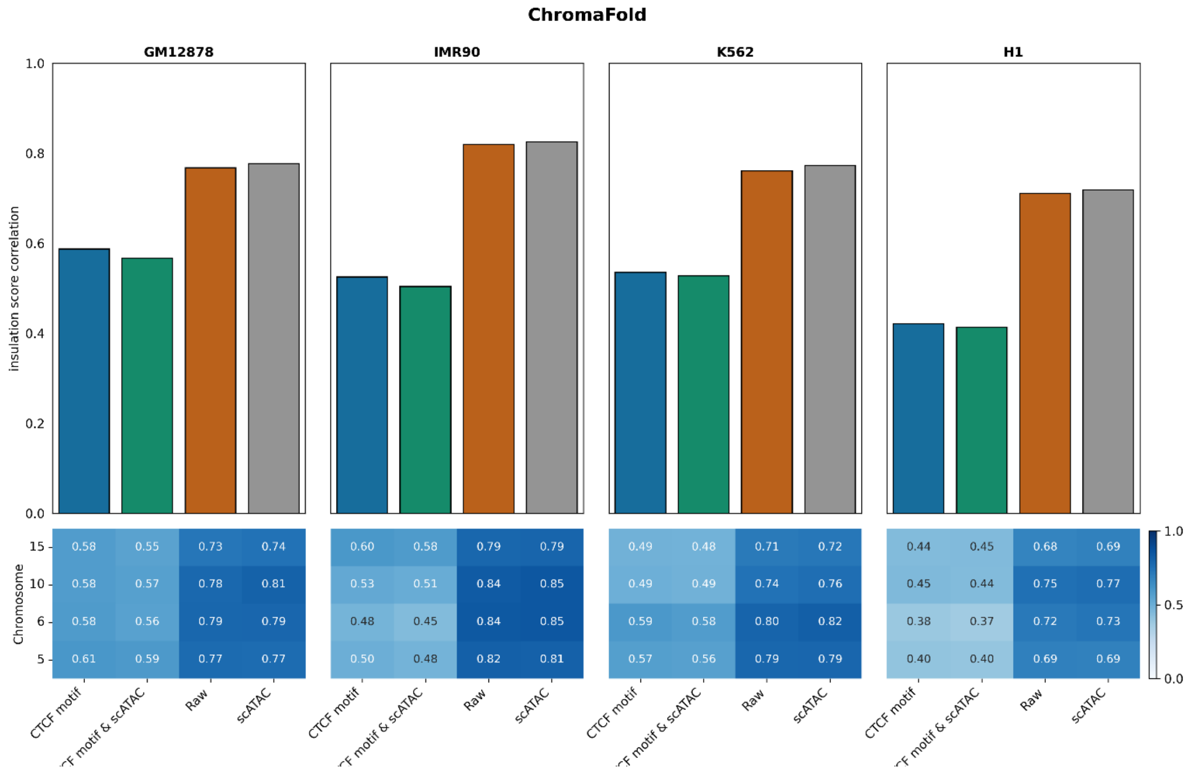


**Supplementary Figure S4.** ChromaFold ablation performance across modalities, cell lines, and chromosomes using insulation score correlation. Top: Bar plot showing ablation results; the x-axis lists the removed modality, and the y-axis shows insulation score correlation. Bottom: Heatmap with chromosomes on the y-axis and removed modalities on the x-axis; each cell's color and value indicate the corresponding performance.

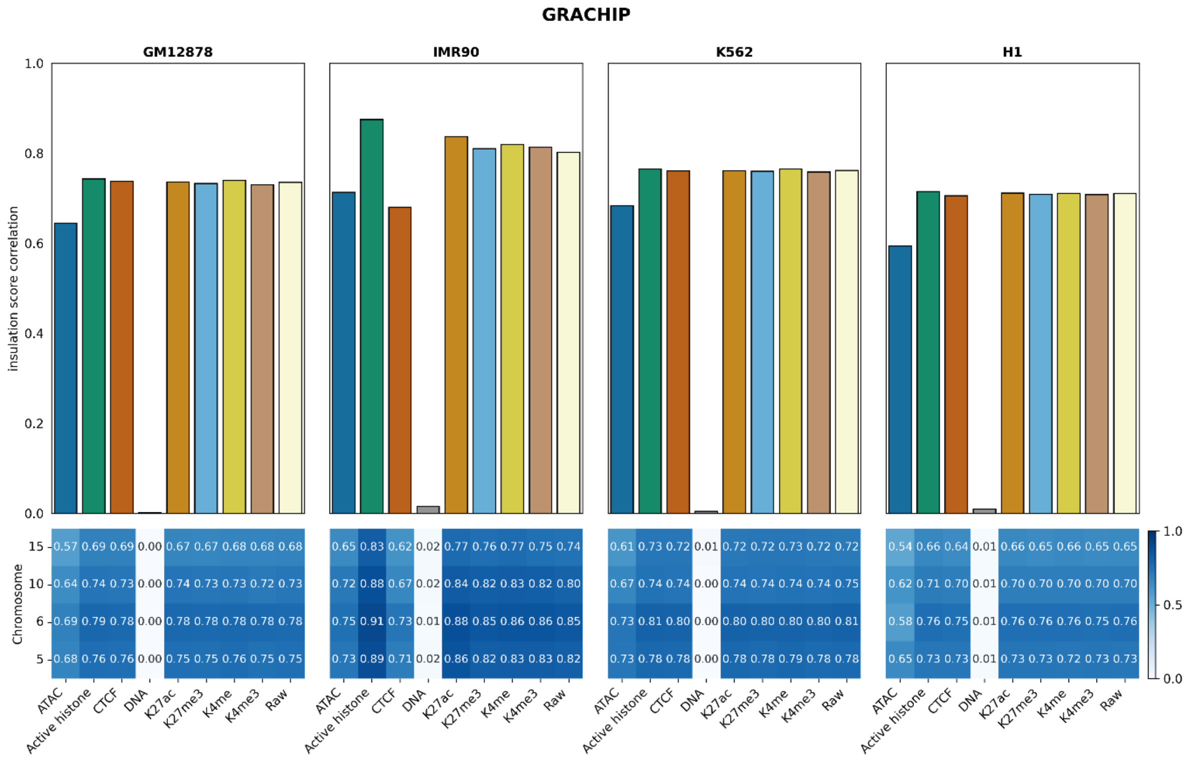


**Supplementary Figure S5.** GRACHIP ablation performance across modalities, cell lines, and chromosomes using insulation score correlation. Top: Bar plot showing ablation results; the x-axis lists the removed modality, and the y-axis shows insulation score correlation. Bottom: Heatmap with chromosomes on the y-axis and removed modalities on the x-axis; each cell's color and value indicate the corresponding performance.

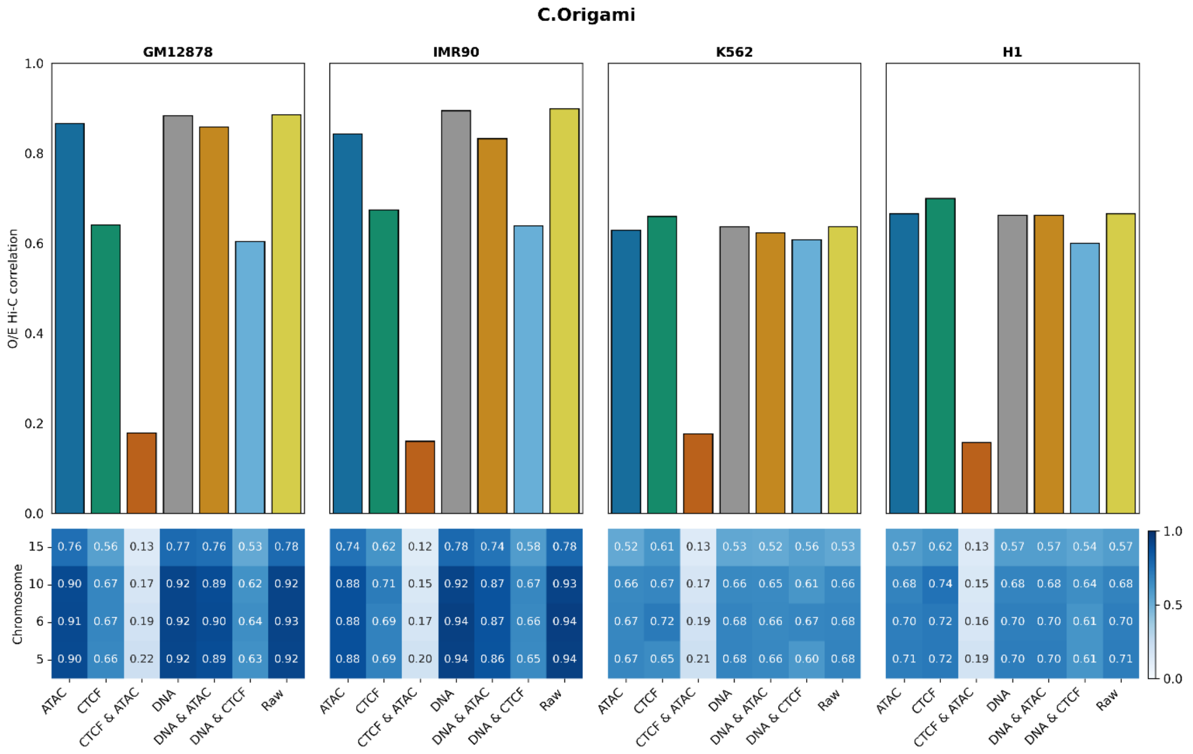


**Supplementary Figure S6**. C.Origami ablation performance across modalities, cell lines, and chromosomes using O/E Hi-C correlation. Top: Bar plot showing ablation results; the x-axis lists the removed modality, and the y-axis shows O/E Hi-C correlation. Bottom: Heatmap with chromosomes on the y-axis and removed modalities on the x-axis; each cell's color and value indicate the corresponding performance.

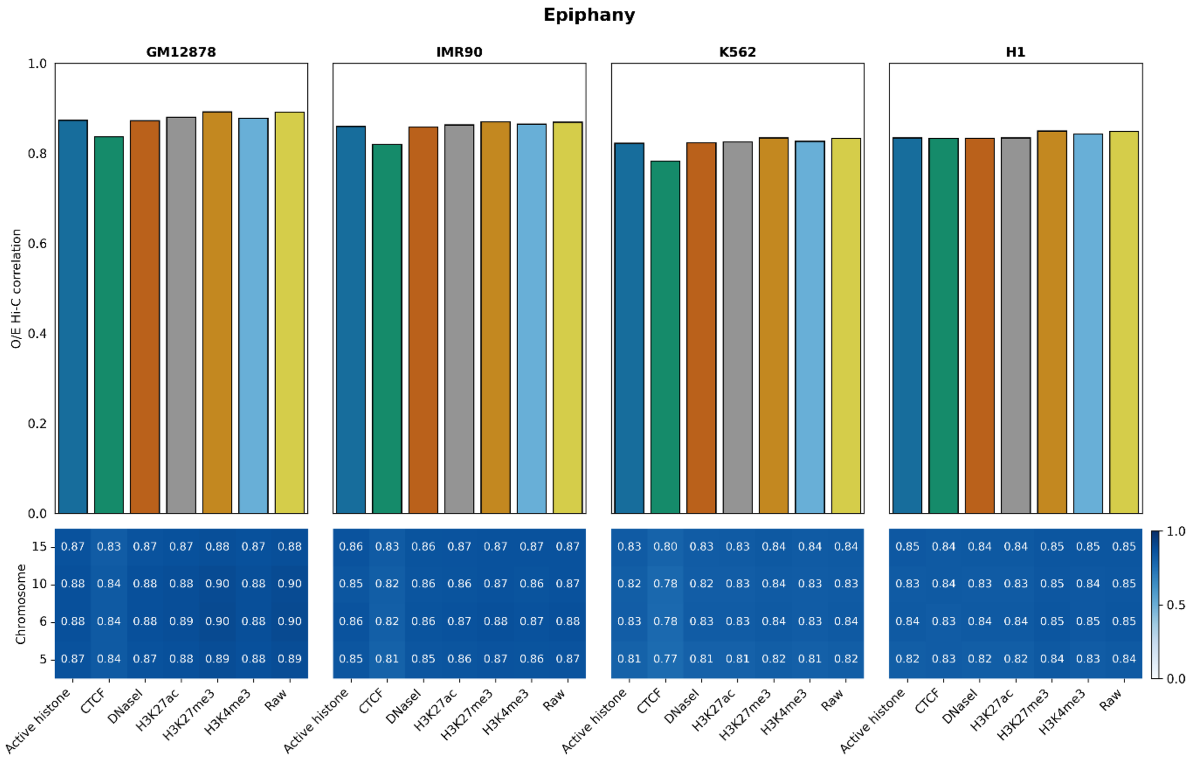


**Supplementary Figure S7.** Epiphany ablation performance across modalities, cell lines, and chromosomes using O/E Hi-C correlation. Top: Bar plot showing ablation results; the x-axis lists the removed modality, and the y-axis shows O/E Hi-C correlation. Bottom: Heatmap with chromosomes on the y-axis and removed modalities on the x-axis; each cell's color and value indicate the corresponding performance.

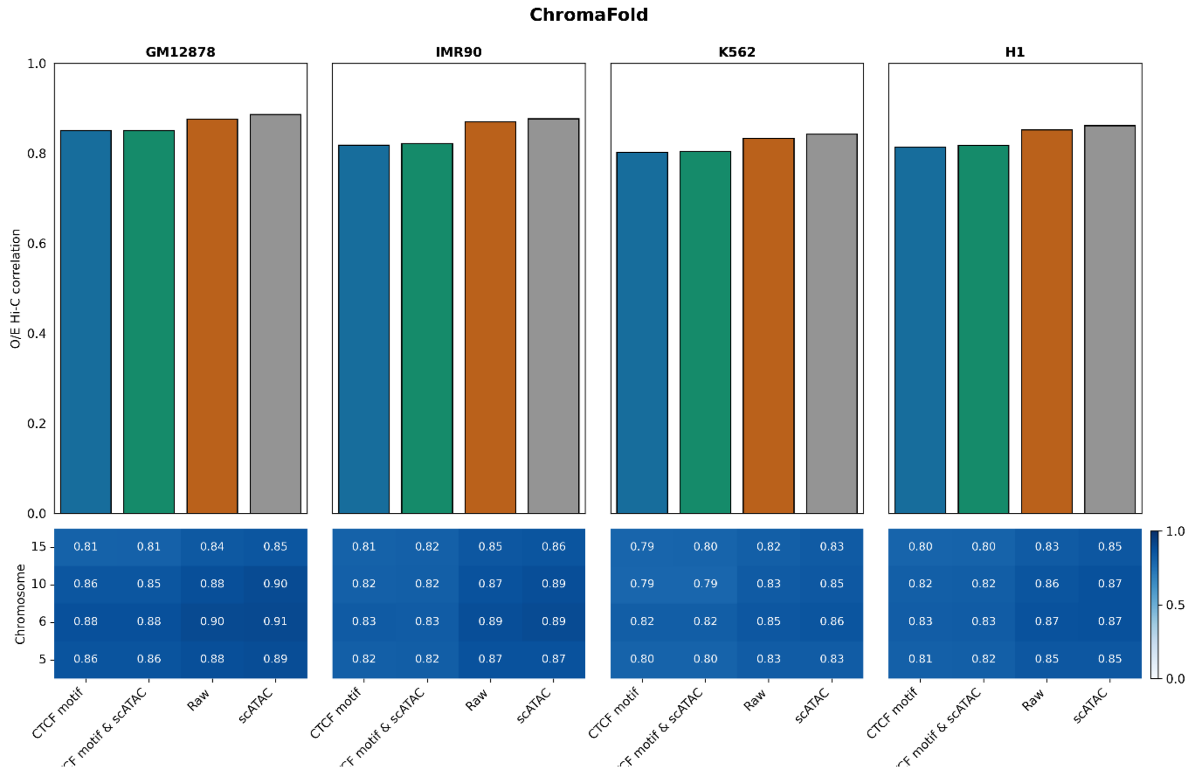


**Supplementary Figure S8.** ChromaFold ablation performance across modalities, cell lines, and chromosomes using O/E Hi-C correlation. Top: Bar plot showing ablation results; the x-axis lists the removed modality, and the y-axis shows O/E Hi-C correlation. Bottom: Heatmap with chromosomes on the y-axis and removed modalities on the x-axis; each cell's color and value indicate the corresponding performance.

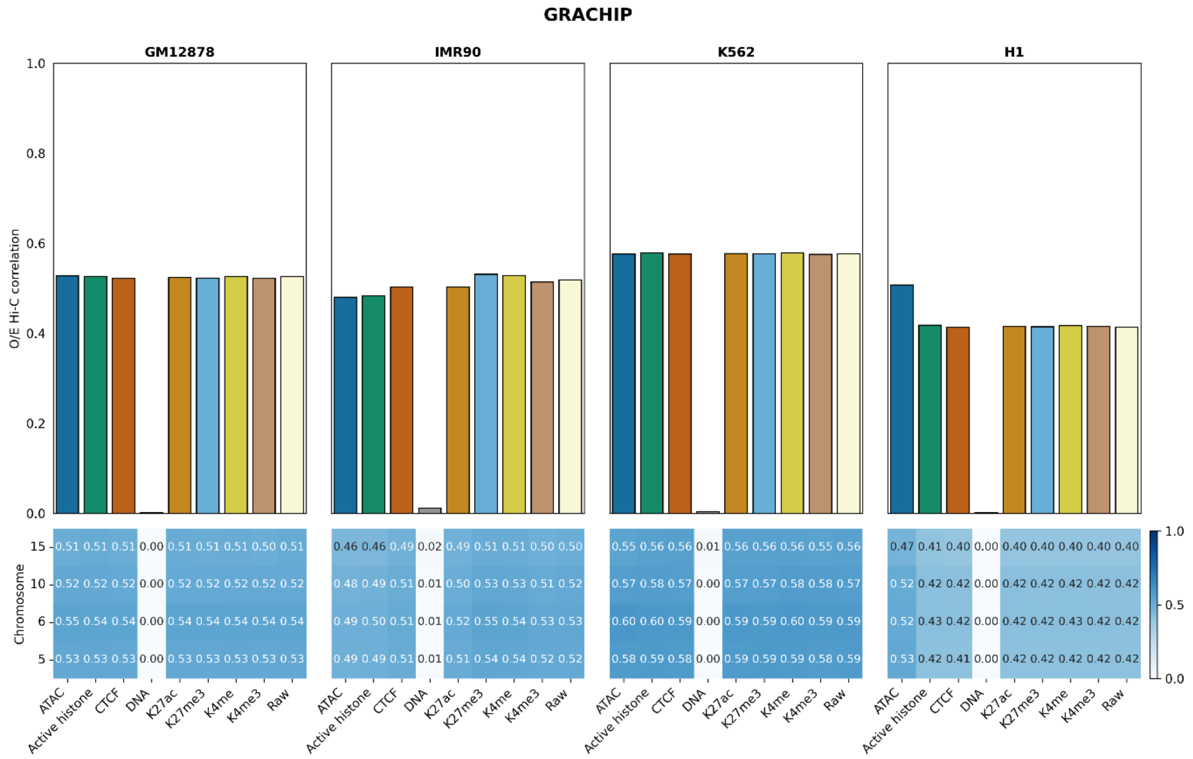


**Supplementary Figure S9.** GRACHIP ablation performance across modalities, cell lines, and chromosomes using O/E Hi-C correlation. Top: Bar plot showing ablation results; the x-axis lists the removed modality, and the y-axis shows O/E Hi-C correlation. Bottom: Heatmap with chromosomes on the y-axis and removed modalities on the x-axis; each cell's color and value indicate the corresponding performance.

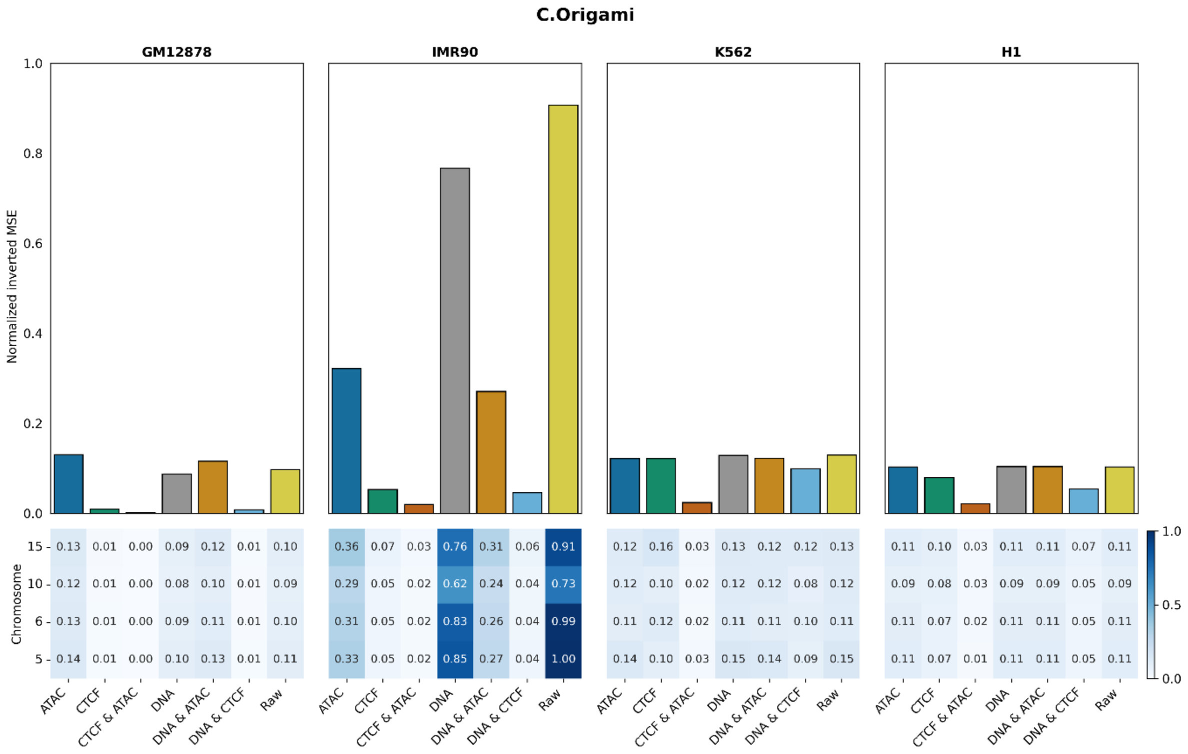


**Supplementary Figure S10.** C.Origami ablation performance across modalities, cell lines, and chromosomes using normalized inverted MSE. Top: Bar plot showing ablation results; the x-axis lists the removed modality, and the y-axis shows normalized inverted MSE. Bottom: Heatmap with chromosomes on the y-axis and removed modalities on the x-axis; each cell's color and value indicate the corresponding performance.

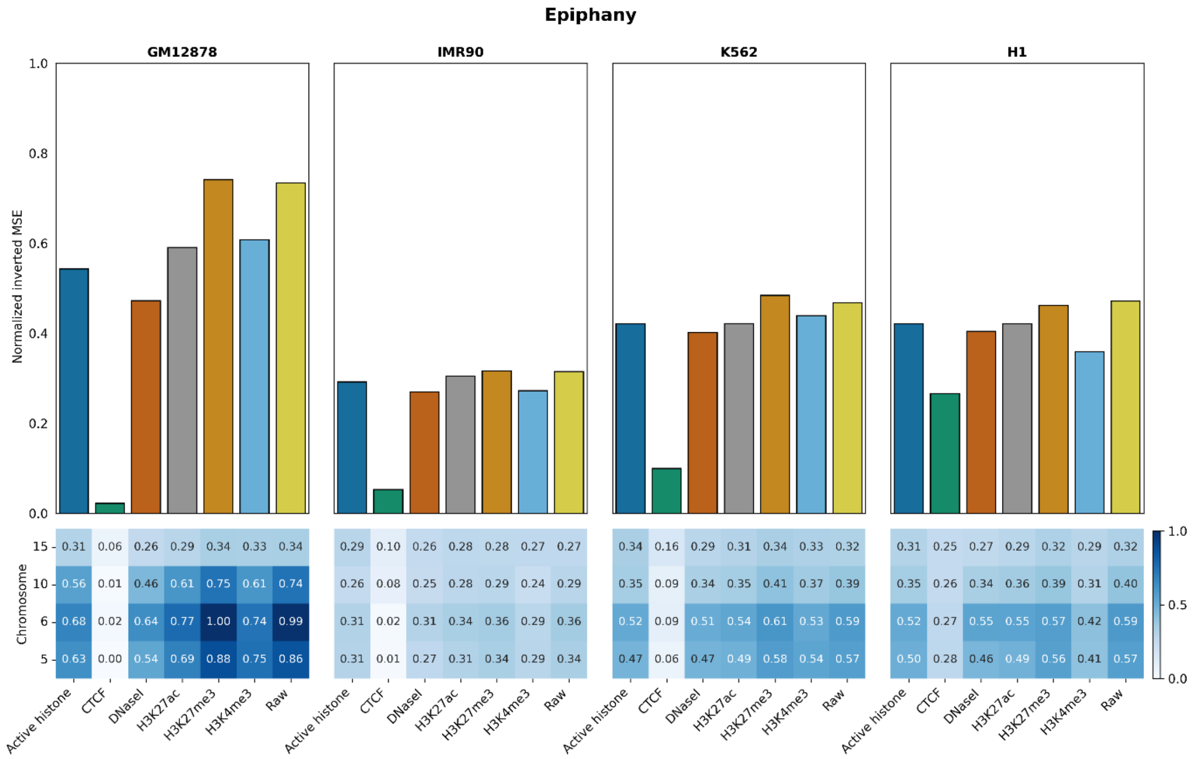


**Supplementary Figure S11.** Epiphany ablation performance across modalities, cell lines, and chromosomes using normalized inverted MSE. Top: Bar plot showing ablation results; the x-axis lists the removed modality, and the y-axis shows normalized inverted MSE. Bottom: Heatmap with chromosomes on the y-axis and removed modalities on the x-axis; each cell's color and value indicate the corresponding performance.

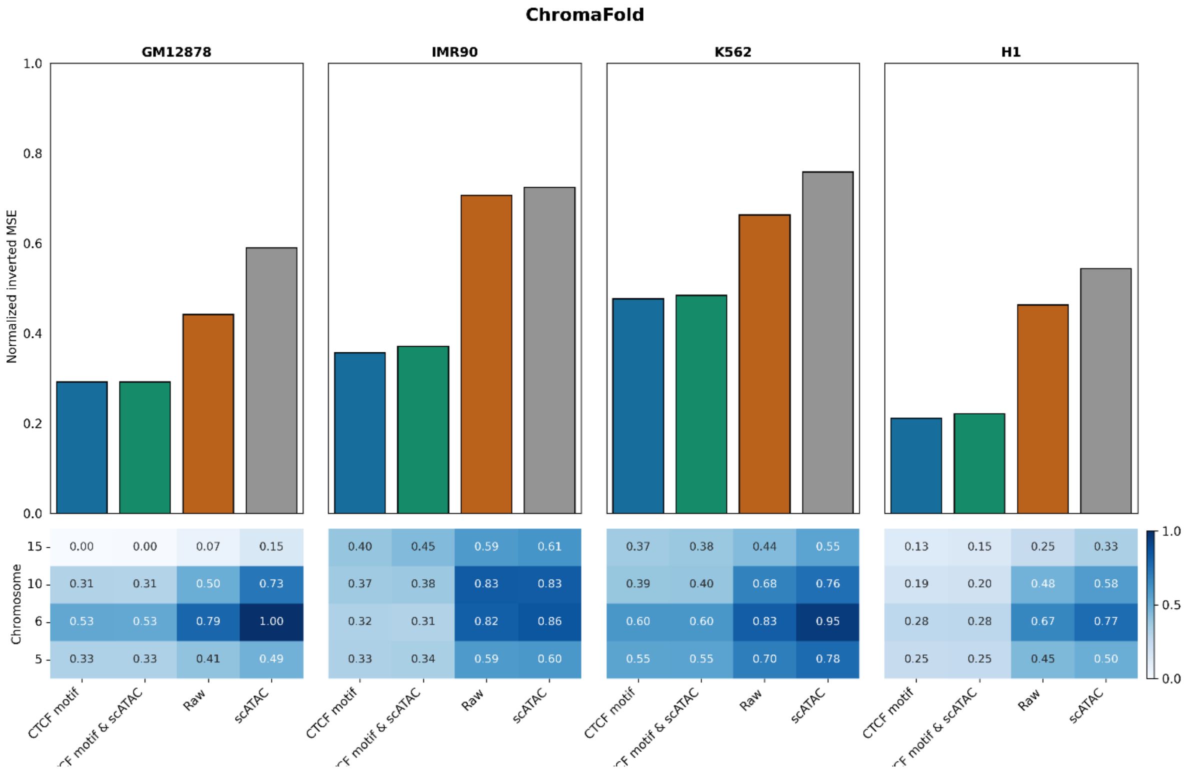


**Supplementary Figure S12.** ChromaFold ablation performance across modalities, cell lines, and chromosomes using normalized inverted MSE. Top: Bar plot showing ablation results; the x-axis lists the removed modality, and the y-axis shows normalized inverted MSE. Bottom: Heatmap with chromosomes on the y-axis and removed modalities on the x-axis; each cell's color and value indicate the corresponding performance.

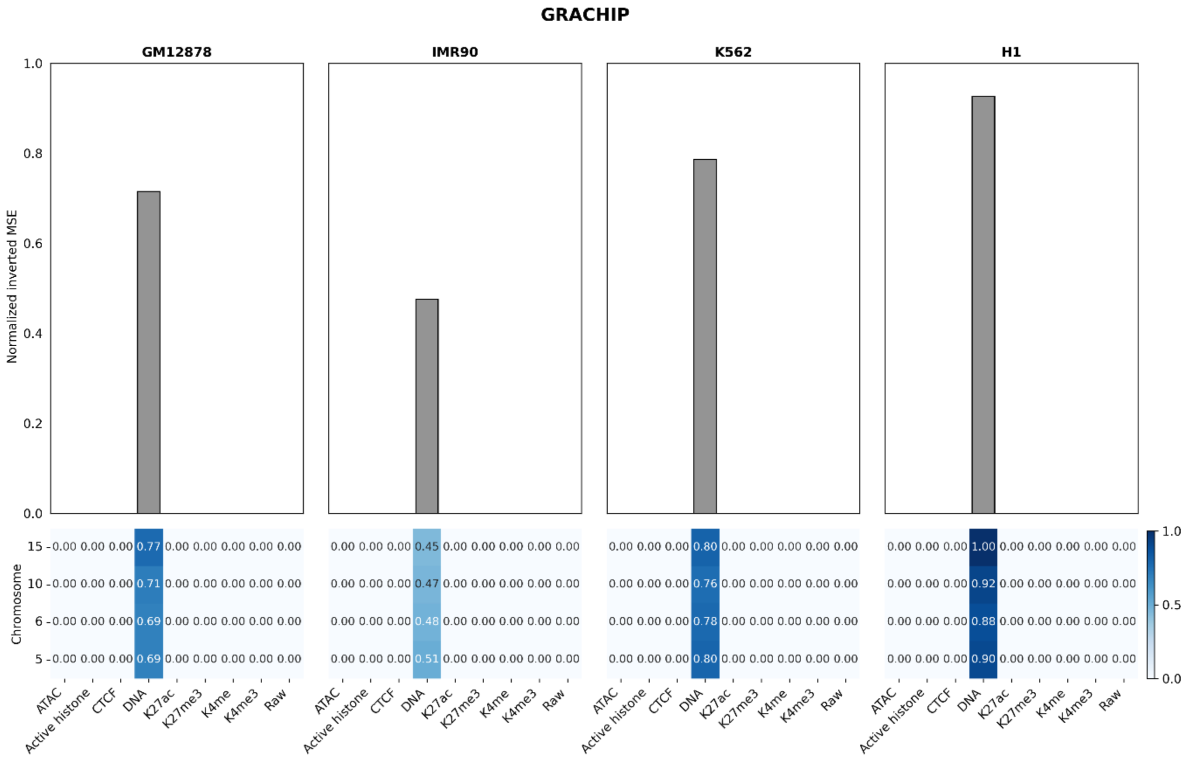


**Supplementary Figure S13.** GRACHIP ablation performance across modalities, cell lines, and chromosomes using normalized inverted MSE. Top: Bar plot showing ablation results; the x-axis lists the removed modality, and the y-axis shows normalized inverted MSE. Bottom: Heatmap with chromosomes on the y-axis and removed modalities on the x-axis; each cell's color and value indicate the corresponding performance.

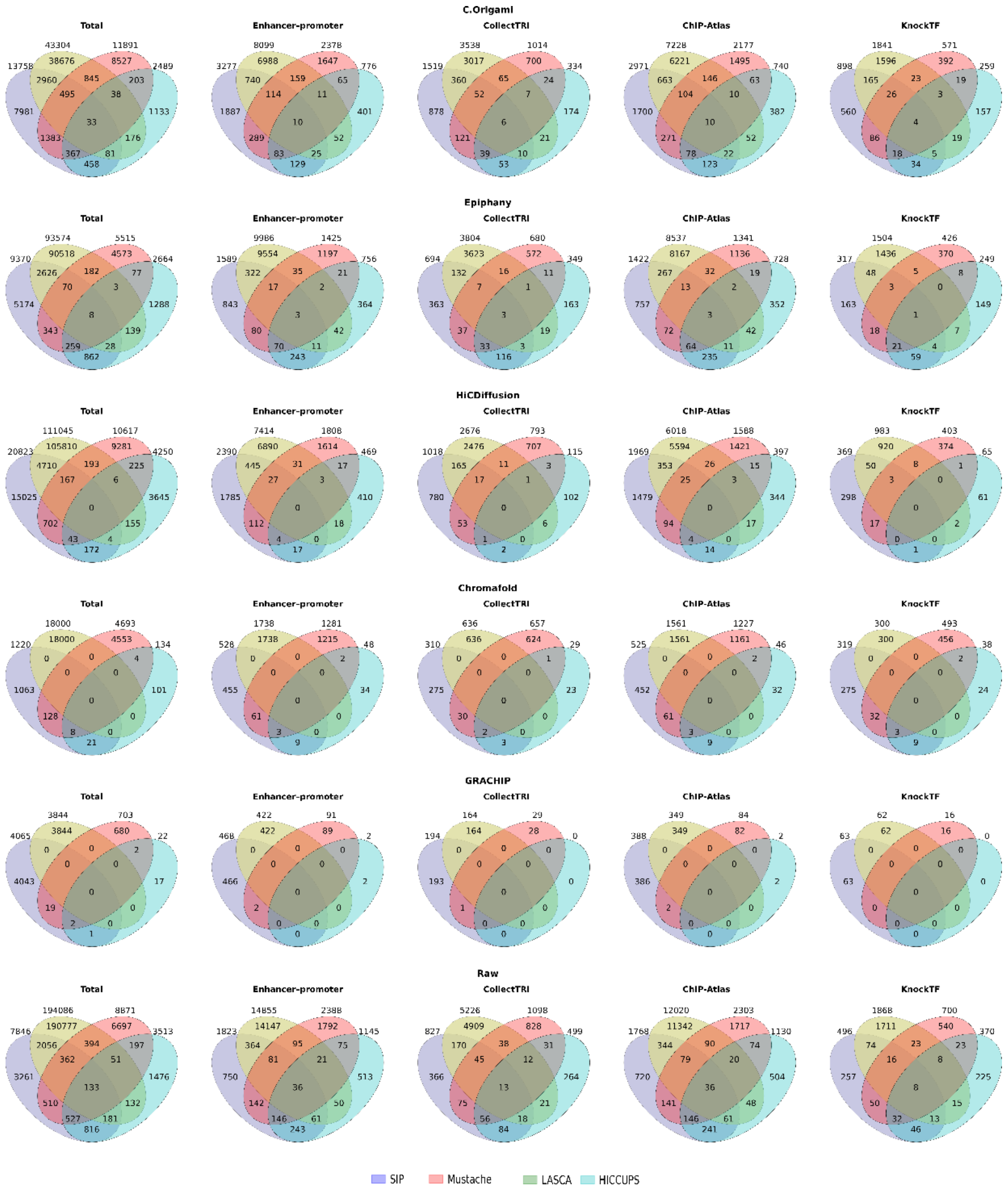


**Supplementary Figure S14.** Loop overlap across four loop-calling tools for each model.

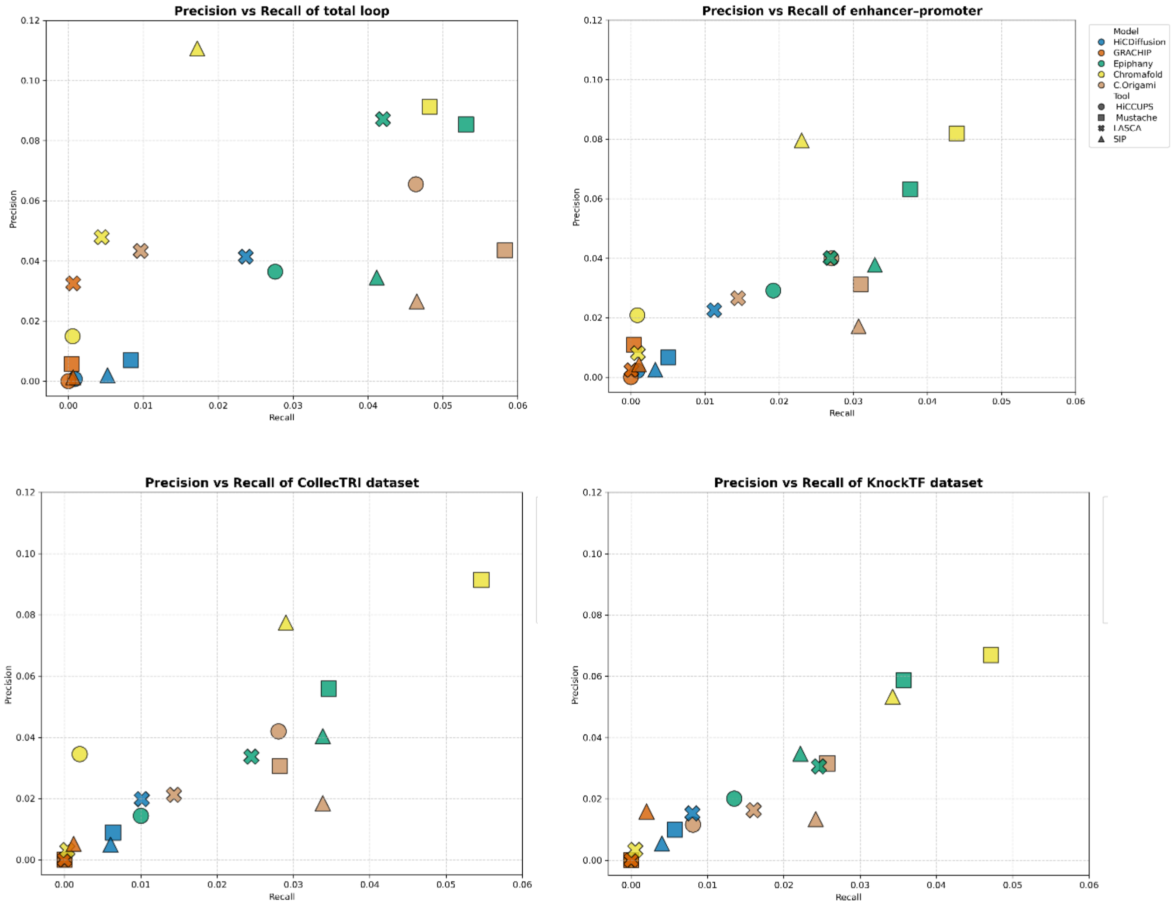


**Supplementary Figure S15.** Precision–recall analysis of total chromatin loops (top-left), loops containing enhancer-promoter pair (top-right), loops supported by CollecTRI dataset (bottom-left), and loops supported by KnockTF dataset (bottom-right). The y-axis indicates precision, and the x-axis indicates recall, using loops derived from the true Hi-C map as the ground truth reference. Each marker represents one combination of a Hi-C prediction model and a loop-calling tool. Marker colour denotes the prediction model, while marker shape denotes the loop caller.
